# RIG-I–MAVS–NOXA axis coordinates antiviral defense and apoptosis during parahenipavirus infection

**DOI:** 10.64898/2026.08.24.746853

**Authors:** Shivani Rajoriya, Divya Misra, Sun Hye Yu, Altanzul Bat Ulzii, Hennisa Hennisa, Tae-Wook Kang, Hye Jin Shin, Yeonsu Oh, Carolina Lopez, Won-Keun Kim

**Affiliations:** Department of Microbiology, College of Medicine, Hallym University, Chuncheon, 24252, Republic of Korea; theMoagen, 32, Doandong-ro 11beon-gil, Seo-gu, Daejeon, 35368, Republic of Korea; Department of Microbiology, School of Medicine, Chungnam National University, Daejeon, 35015, Republic of Korea; College of Veterinary Medicine and Institute of Veterinary Science, Kangwon National University, Chuncheon, 24341, Republic of Korea; Department of Molecular Microbiology and Center for Women Infectious Disease Research, Washington University School of Medicine in St. Louis, Missouri, 63110, United States of America; Institute of Medical Research, College of Medicine, Hallym University, Chuncheon, 24252, Republic of Korea

**Keywords:** Gamak virus, Parahenipavirus, RIG-I, MAVS, Type I interferon, NOXA, Apoptosis, Innate immunity

## Abstract

The Gamak virus (GAKV) is a recently identified shrew-borne paramyxovirus belonging to the genus *Parahenipavirus*, which also includes the zoonotic Langya virus (LayV). Despite the growing recognition of shrew-associated paramyxoviruses, the host pathways that detect infection and regulate antiviral responses remain poorly understood. In this study, we characterized host responses to GAKV infection using integrated *in vitro* and *in vivo* approaches. GAKV infection induced robust innate immune responses in A549 cells, characterized by activation of interferon regulatory factor 3 (IRF3) and signal transducer and activator of transcription 1 (STAT1), together with induction of type I interferon (IFN) and interferon-stimulated genes (ISGs). Transcriptomic analysis further revealed coordinated enrichment of antiviral and intrinsic apoptosis-associated pathways, suggesting a link between innate immune signaling and apoptosis during GAKV infection. Genetic analyses identified retinoic acid-inducible gene I (RIG-I) and mitochondrial antiviral signaling protein (MAVS) as essential mediators of antiviral signaling and apoptosis during GAKV infection. Furthermore, disruption of type I IFN-STAT1 signaling attenuated apoptosis. NOXA knockdown reduced apoptosis and enhanced viral replication, identifying NOXA as a downstream effector linking innate immune activation to apoptosis. Consistent with these *in vitro* findings, intranasal GAKV infection in six-week-old female wild-type BALB/c mice was associated with lung-restricted viral RNA detection and induction of antiviral responses without overt disease. Together, these findings identify a RIG-I-MAVS-IFN-NOXA signaling axis that integrates antiviral and apoptotic responses during GAKV infection, providing a mechanistic framework for understanding host defense against parahenipaviruses.

**Impact Statement:** The recently identified Gamak virus (GAKV) is a member of the genus *Parahenipavirus*; however, how host cells recognize GAKV infection and initiate antiviral defenses remains unknown. We found that GAKV is primarily detected by the innate immune sensor retinoic acid-inducible gene I (RIG-I), which activates antiviral signaling through the adaptor protein mitochondrial antiviral signaling protein (MAVS) and initiates type I interferon (IFN) responses. In addition, this pathway promotes programmed cell death (apoptosis) through downstream induction of NOXA. GAKV infection induced antiviral responses in the lungs of mice without causing overt disease. Together, our findings establish a framework for investigating immune responses to newly identified parahenipaviruses.

## Introduction

The *Paramyxoviridae* family includes multiple human and animal pathogens, including the measles, Hendra (HeV), and Nipah viruses (NiV) [1]. Paramyxoviruses are enveloped, non-segmented, negative-sense, single-stranded RNA viruses. During infection in the host cell, paramyxovirus transcription proceeds sequentially from the 3′ end to the 5′ end of the genome through a start-stop mechanism, resulting in a transcriptional gradient in which genes near the 3′ end are expressed at higher levels than downstream genes [2].

Recent surveillance studies have uncovered extensive paramyxovirus diversity in wildlife reservoirs, leading to the discovery of multiple novel viruses with poorly understood replication characteristics and host-cell interactions [3–6]. For instance, Gamak virus (GAKV) is a recently identified shrew-borne paramyxovirus isolated from *Crocidura lasiura* in the Republic of Korea, classified under the genus *Parahenipavirus* [7]. The genus also includes Langya virus (LayV), which has been associated with human infections, underscoring the potential public health relevance of parahenipaviruses [8]. Although members of the genus *Parahenipavirus* have attracted increasing attention following their recent discovery, the host pathways that detect parahenipavirus infection and coordinate antiviral responses remain poorly understood. Specifically, the mechanisms that sense GAKV infection and initiate antiviral defenses have not been defined, and *in vivo* host responses to GAKV infection remain unknown.

Viral RNA in host cells is detected by cytosolic pattern recognition receptors (PRRs), thereby activating innate immune defenses [9]. Among these PRRs, retinoic acid-inducible gene I (RIG-I) and melanoma differentiation associated factor 5 (MDA5) play central roles in sensing signatures contained within viral RNA [10]. Upon activation, RLRs signal through the mitochondrial antiviral signaling protein (MAVS), triggering phosphorylation of interferon regulatory factor 3 (IRF3) and subsequent induction of type I interferon (IFN) expression [11]. IFN-β binds to IFNAR1 and IFNAR2, which are expressed on nucleated cells. This activates the JAK-STAT signaling pathway, leading to the formation of the interferon-stimulated gene factor 3 (ISGF3) complex, consisting of STAT1, STAT2, and IRF9, which drives transcription of interferon-stimulated genes (ISGs). However, the relative contribution of individual RLRs to antiviral sensing varies between paramyxoviruses and host cell types [12, 13]. In addition to IFN responses, viral infections induce chemokines that recruit and activate immune cells, thereby contributing to antiviral defense and shaping inflammatory responses [14].

Innate immune signaling pathways can influence cell fate decisions by inducing pro-apoptotic genes and activating programmed cell death pathways [15]. Apoptosis is an important antiviral defense mechanism that limits viral replication and dissemination [16]. The intrinsic apoptotic pathway is regulated by B cell lymphoma 2 (Bcl-2) family proteins, including BCL- 2 homology 3 (BH3)-only proteins, which promote mitochondrial outer membrane permeabilization and activation of caspase-9. In contrast, the extrinsic pathway is initiated through death receptor signaling and activation of caspase-8. Nevertheless, both pathways converge upon activation of executioner caspases, including caspase-3 [17]. Although apoptosis has been reported during infection with multiple paramyxoviruses, the molecular mechanisms linking innate immune sensing to apoptotic responses remain poorly understood, particularly in the newly identified shrew-borne paramyxoviruses [2, 18–20].

In this study, we investigated GAKV replication and host responses using complementary *in vitro* and *in vivo* approaches. We aimed to define the host pathways that sense GAKV infection and coordinate antiviral and apoptotic responses. We used an immunocompetent BALB/c mouse model to investigate the *in vivo* host response to GAKV infection, including tissue tropism and antiviral responses. Our findings identify a RIG-I-MAVS-NOXA-dependent host defense pathway and provide the first comprehensive characterization of host responses to GAKV infection.

## Materials and Methods

### Ethics statement

*In vitro* experiments were conducted at Biosafety Level-2 in accordance with the guidelines and protocols of the Hallym University Institutional Biosafety Committee. All animal experiments were conducted in the Animal Biosafety Level-2 (ABSL-2) facility at the Research Institute of Medical Bio-Convergence of Hallym University. The animal experiments were approved by the Institutional Animal Care and Use Committee (IACUC) of Hallym University (Approval No. HallymR1 2024-11). Anesthesia was induced in the mice via an intraperitoneal injection of 2,2,2-tribromoethanol dissolved in tert-amyl alcohol (Sigma-Aldrich, T48402) in accordance with the IACUC-approved protocols. At the designated experimental time points, mice were euthanized by cervical dislocation for tissue collection.

### Cell culture

A549 (ATCC, CCL-185), Huh7 (Korean Cell Line Bank, 60104), and Vero E6 (ATCC, CRL-1586) cells were maintained in Dulbecco’s modified Eagle’s medium (DMEM; Gibco, 11995065) supplemented with 10% Fetal Bovine Serum (Gibco, A5669701), 1% of 1 M HEPES (15630-080, Gibco), and 1% antibiotic-antimycotic (15240062, Gibco). All cell lines were cultured at 37 °C and 5% CO_2_ in a humidified incubator. A549 CRISPR control and STAT1-knockout (STAT1 KO) cell lines were kindly provided by Dr. Susan R. Weiss (University of Pennsylvania) and were previously characterized [21].

### Animals and GAKV infection

30 six-week-old female BALB/c mice were purchased from Central Lab Animal Inc., Republic of Korea, and housed in a temperature- and humidity-controlled room (23 °C and 50%-55% relative humidity) under a 12-h light/dark cycle. Following a one-week acclimation period with ad libitum access to standard rodent chow and water, mice were randomly assigned to experimental groups. The animals were divided into two groups: mock controls (n = 15) and GAKV-infected mice (n = 15). Each group was further subdivided into three cohorts for tissue collection at 1-, 3-, and 5-days post-infection (dpi) (n = 5 per group per time point). Before infection, the mice were anesthetized and intranasally administered GAKV (5 × 10^4^ PFU per mouse) by carefully pipetting the inoculum onto the nostrils. Following infection, the animals were monitored daily for clinical signs and body weight changes. At the designated experimental time points, mouse tissues were collected aseptically, immediately homogenized in TRIzol reagent, and stored at -80 °C until RNA extraction and RT-qPCR analysis. Mock control mice were administered sterile phosphate-buffered saline (PBS).

### Plaque assay

GAKV titration was performed using Vero E6 cells seeded at 1 × 10^6^ cells per well in six-well plates. The cell monolayers were washed with PBS, and 10-fold serial dilutions of the virus were added to each well. The plates were incubated for 2 h at 37 °C to allow viral adsorption. Subsequently, the overlay medium, prepared in 2Χ DMEM/F-12 supplemented with 2% Oxoid agar (LP0028, Thermo Scientific), was added to each well, and cells were incubated for six days at 37 °C in a 5% CO_2_ incubator. The fixed cell monolayers were stained with 0.1% crystal violet solution. Viral titers were calculated and expressed as plaque-forming units per mL (PFU/mL).

### Confocal immunostaining

A total of 5 × 10^4^ A549 cells were seeded on lysine-coated glass coverslips in 12-well plates. On the following day, the cells were infected with GAKV at the indicated multiplicity of infection (MOI) and incubated at 37 °C in a 5% CO_2_ incubator for 48 h. For the positive control in the apoptosis assay, the cells were incubated with staurosporine (Cell Signaling Technology CST #9953) for 3 h in the infection medium before fixation. Following fixation, the cells were permeabilized and stained for 2 h with primary antibodies against GAKV-N (Abclonal Korea, 1:12,800, rabbit) and cleaved caspase-3 (Cell Signaling Technology CST #9661S, 1:400, rabbit). Subsequently, the cells were washed and stained with fluorophore-conjugated secondary antibodies: Alexa Fluor 488 (Invitrogen, A-11008, goat anti-rabbit, 1:500) and Alexa Fluor 568 (Invitrogen, A-11011, goat anti-rabbit, 1:500). Nuclei were then counterstained with Hoechst 33342 (Invitrogen, 62249, 1:500). Coverslips were mounted onto glass slides using the Dako Fluorescence Mounting Medium (Agilent Technologies, S3023).

Apopxin Green staining was performed according to the manufacturer’s instructions (ab176749; Abcam). Briefly, Apopxin Green was diluted 1:100 in assay buffer and incubated with cells for 45 min at room temperature. Cells were then washed with assay buffer, fixed, and stained for GAKV N protein by immunofluorescence. Images were acquired using a Carl Zeiss super-resolution confocal laser-scanning microscope (LSM900 with Airyscan2). Images were merged and processed using the ZEN Blue software (Carl Zeiss Microscopy GmbH, v3.4.91.00000). Further analyses of the images were performed using the ImageJ software (version 1.52; NIH, USA).

### Western blotting

A549 cells were seeded at 0.3 × 10^6^ cells per well in six-well plates and cultured overnight before infection. Following the indicated time points, cells were harvested and lysed using RIPA buffer (CST, #9806) supplemented with Protease/Phosphatase Inhibitor Cocktail (CST, #5872) for western blot analysis. Protein concentrations were determined by the bicinchoninic acid (BCA) assay. Equal amounts of protein (45 μg per lane) were separated by SDS-PAGE, with the T&I™ ACCU Prestained Protein Marker (PJM-0605N) used as the molecular weight marker. Transfer membrane (Immobilon-P IPVH00010) were incubated overnight at 4 °C with primary antibodies against GAKV N (custom-made polyclonal antibody, Abclonal Korea, 1:1000, rabbit), phospho IRF3 (CST #4947, 1:1000, rabbit), IRF3 (CST #4302, 1:1000, rabbit), pIκBα (CST #9246, 1:1000, mouse), IκBα (CST #4814, 1:1000, mouse), pSTAT1 (CST #9167, 1:1000, rabbit), STAT1 (CST #14994, 1:1000, rabbit), OASL (Invitrogen, #MA5-50518, 1:1000, rabbit), MAVS (CST #3993, 1:1000, rabbit), MDA5 (CST #5321, 1:1000, rabbit), RIG-I (CST #3743, 1:1000, rabbit), PMAIP1/NOXA (CST #14766, 1:1000, rabbit), cleaved caspase-8 (CST #98134, 1:1000, rabbit), cleaved caspase-9 (CST #20750, 1:1000, rabbit), cleaved caspase-3 (CST #9661, 1:1000, rabbit), pro caspase-9 (CST #9502, 1:1000, rabbit), pro caspase-8 (CST #4790, 1:1000, rabbit), pro caspase-3 (CST #9662, 1:1000, rabbit) and GAPDH (Sigma-Aldrich G9545, 1:10,000, rabbit). The membranes were then incubated with horseradish peroxidase-conjugated secondary antibodies (Jackson ImmunoResearch 111-035-144, 1:10,000, anti-rabbit; Jackson ImmunoResearch 715-035-150, 1:10,000, anti-mouse) for 1 h at room temperature, followed by chemiluminescence detection. Further analyses of the images were performed using the ImageJ software (version 1.52; NIH, USA).

### RNA Extraction, reverse-transcription polymerase chain reaction (RT-PCR), and quantitative polymerase chain reaction (qPCR)

Cell lysates were dissolved in 1 mL of TRIzol reagent, and total RNA was extracted from uninfected and virus-infected cells according to the manufacturer’s guidelines. RNA was quantified using a NanoDrop 2000 spectrophotometer (Thermo Fisher Scientific). cDNA was prepared from 1 µg of RNA using a High-Capacity RNA-to-cDNA Kit (4387406; Applied Biosystems). Quantitative polymerase chain reaction (qPCR) was performed using the SYBR Green PCR Master Mix (Applied Biosystems, 4367659) on an Applied Biosystems QuantStudio 5 (Thermo Fisher Scientific). The reactions were conducted in 10 µL volumes in a 96-well plate, with thermal cycling conditions starting with an initial denaturation at 95 °C for 30 s, followed by 40 cycles of 95 °C for 15 s and 60 °C for 60 s. Details of the RT-qPCR primer sequences for the GAKV *N* gene are listed in Table 1, human target genes are listed in Table 2, and mouse target genes are listed in Table 3.

**Table 1:**
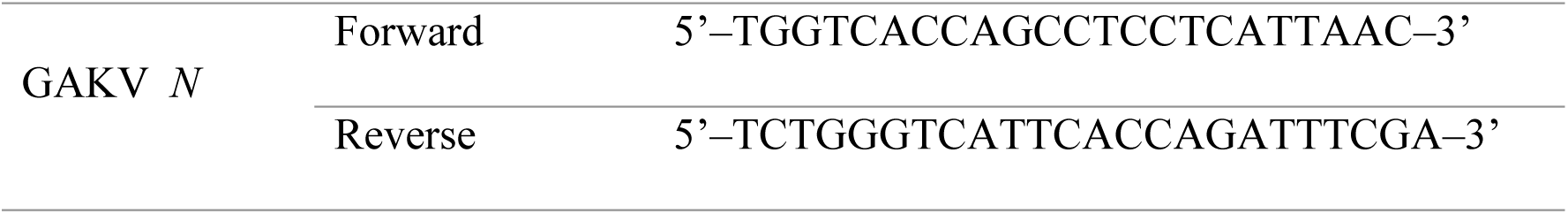
Primers for RT-qPCR analysis of the GAKV *N* gene.

**Table 2:**
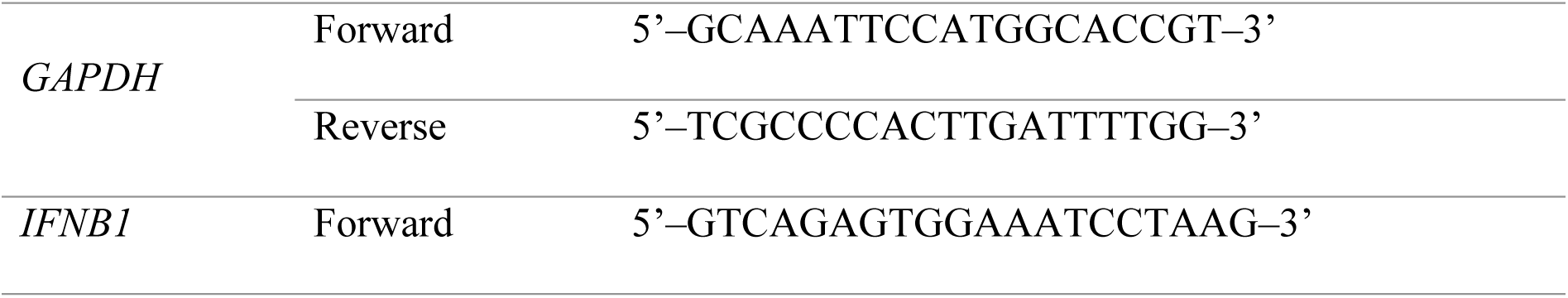
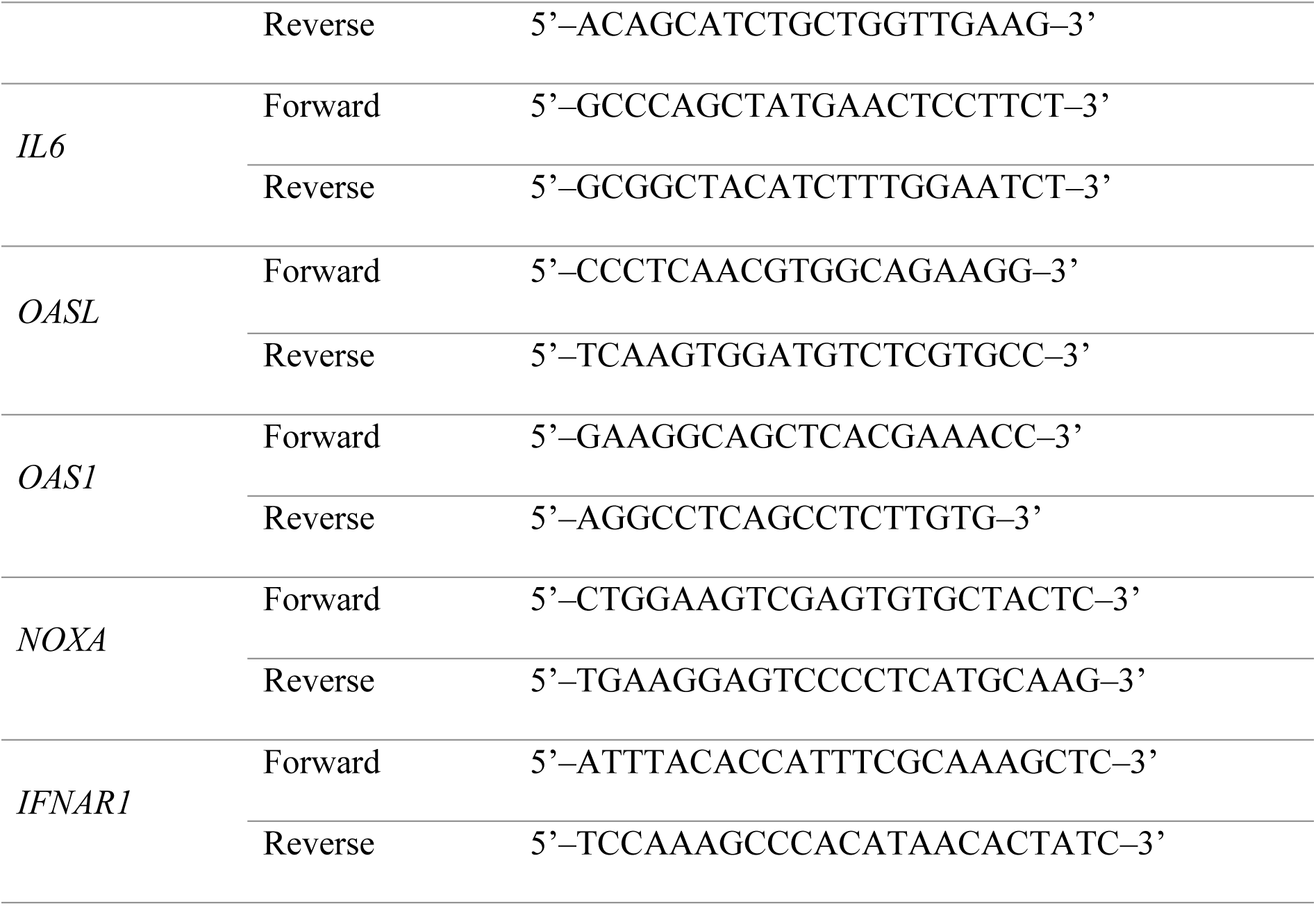
Primers for RT-qPCR analysis of the human genes.

**Table 3:**
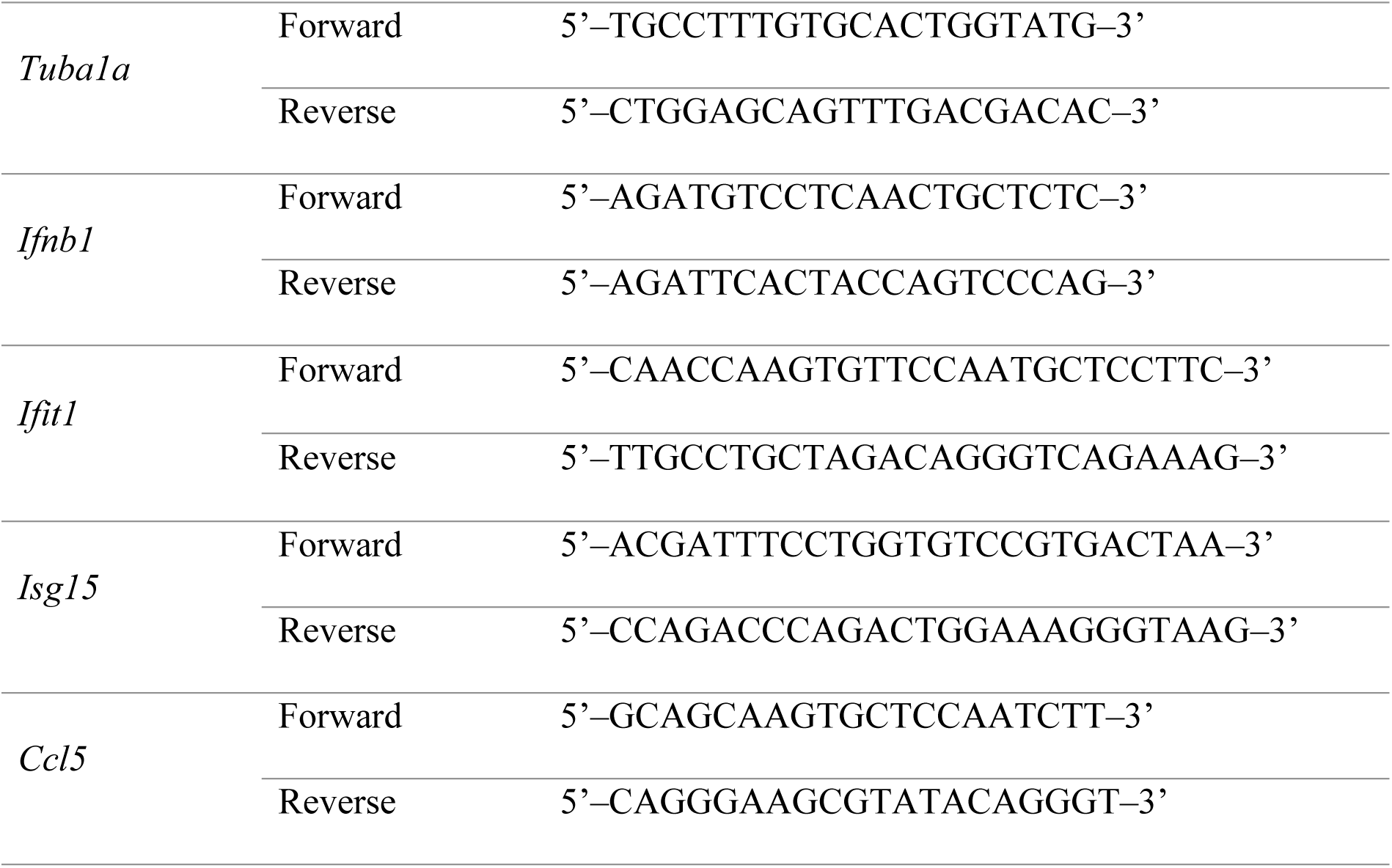
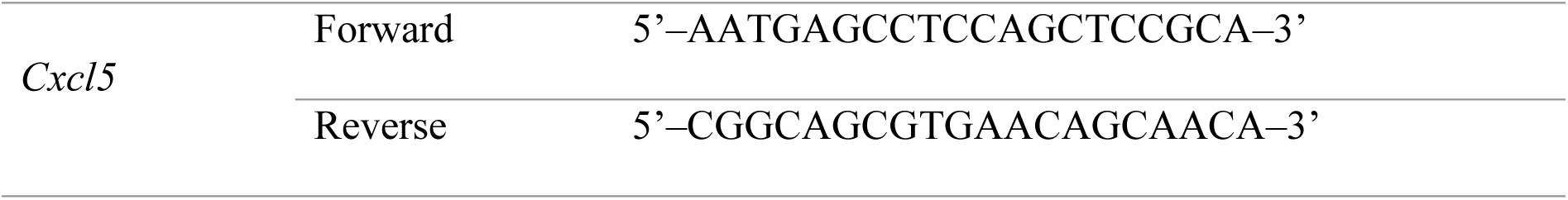
Primers for RT-qPCR analysis of mouse genes.

### Generation and validation of CRISPR-Cas9 knockout A549 cells

CRISPR-Cas9 KO cell lines were generated using predesigned sgRNAs listed in Table 4 (Thermo Fisher Scientific) and TrueCut Cas9 Protein v2 (Invitrogen, A36496). The sgRNA was mixed with the Cas9 protein according to the manufacturer’s instructions to generate CRISPR ribonucleoprotein complexes. Briefly, sgRNA and Cas9 protein were combined in Opti-MEM and incubated at room temperature for approximately 10-15 min to form Cas9-sgRNA ribonucleoprotein (RNP) complexes. The Cas9-sgRNA RNP complexes were introduced into the A549 cells using Lipofectamine CRISPRMAX (Invitrogen, CMAX00008). Single-cell clones were isolated by limiting dilution, expanded, and screened for successful gene disruption. Genomic DNA was extracted from candidate clones, and the target locus was amplified by PCR using the primers listed in Table 5. The PCR products were subjected to Sanger sequencing to confirm the presence of insertions or deletions (indels) at the sgRNA target site. Functional validation of KO clones was further assessed by western blot analysis following stimulation with IFN-β (PBL assay science, 11415) for 14 h, in which the absence of the target protein or downstream signaling confirmed successful gene disruption (S4 Fig).

**Table 4.**
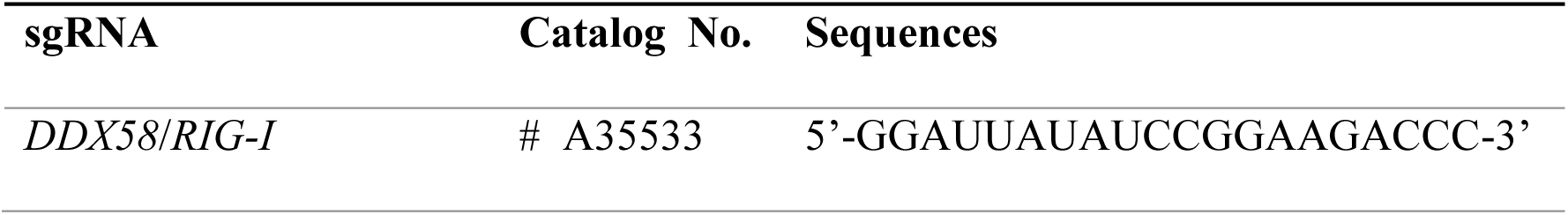
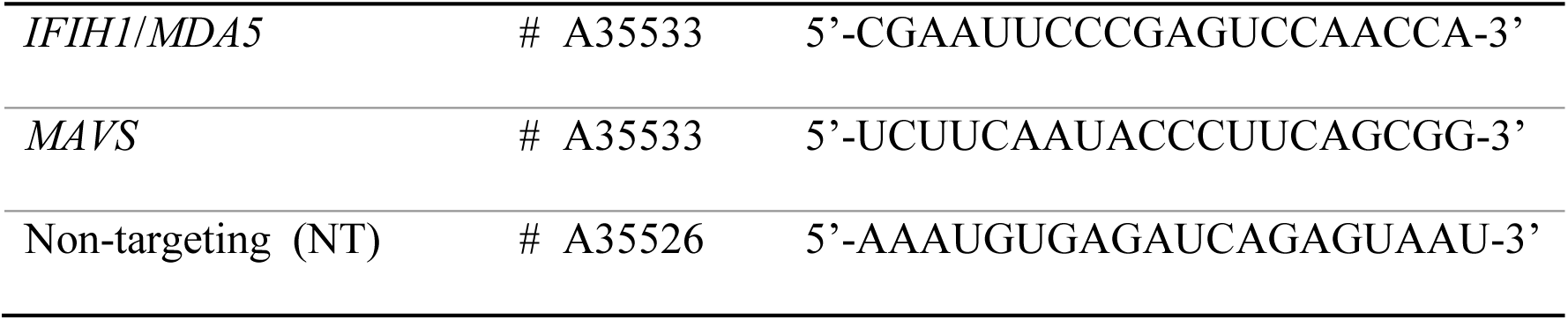
sgRNA sequences for the generation of RIG-I, MDA5, and MAVS knockout A549 cell lines.

**Table 5.**
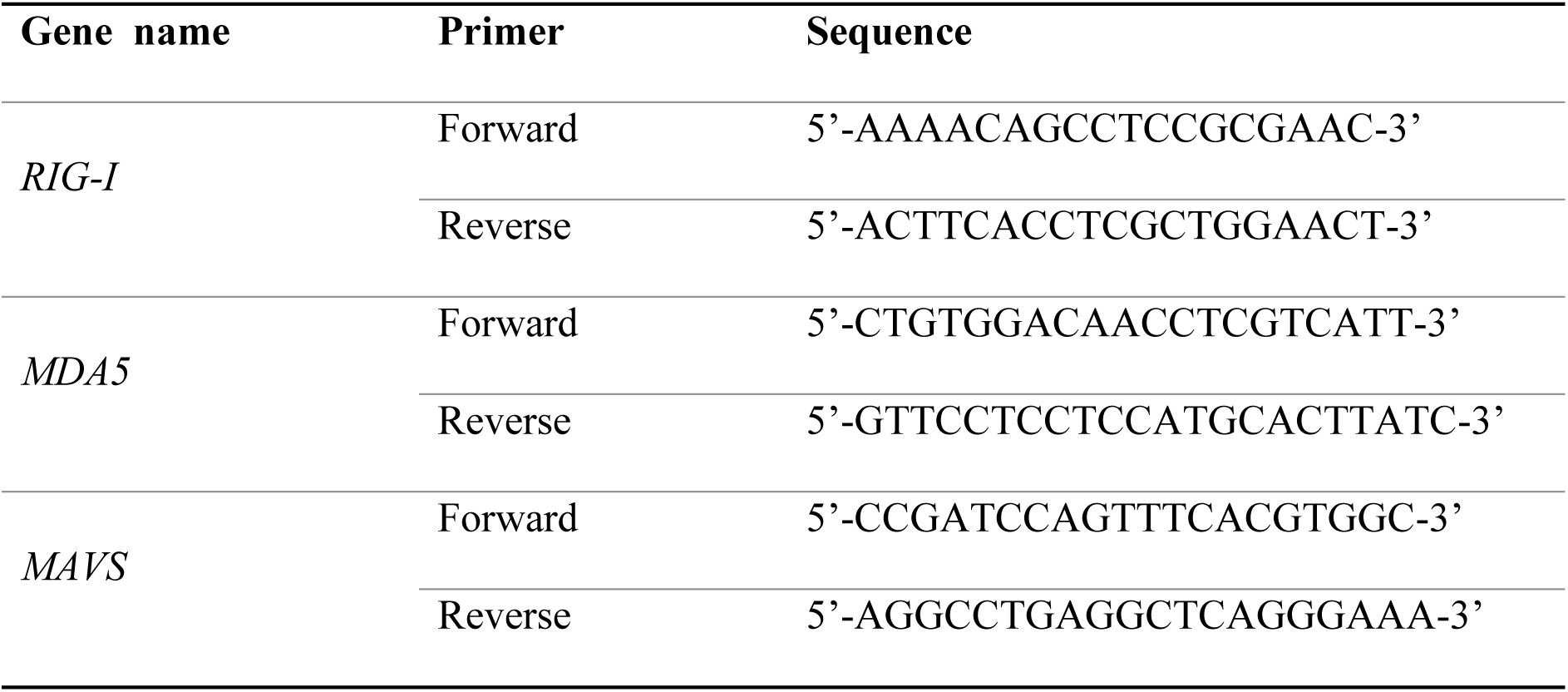
Primer sequences used for validation of RIG-I, MDA5, and MAVS KO A549 cell lines by Sanger sequencing.

### Cell cytotoxicity

Cell death was quantified by measuring the release of lactate dehydrogenase (LDH) into the culture supernatant using the Cytotoxicity LDH Assay Kit-WST (Dojindo, CK12-01), according to the manufacturer’s instructions. Briefly, equal volumes of the supernatants were transferred to 96-well plates. The LDH working solution was prepared according to the manufacturer’s instructions and added to each well. The plates were then incubated at room temperature for 30 min. Absorbance was measured using a microplate reader at 490 nm with a reference wavelength of 490 nm.

### siRNA transfection

siRNA transfection was performed using the DharmaFECT 1 transfection reagent (Horizon Discovery, T-2001-02) according to the manufacturer’s instructions. The siRNA sequences used are listed in Table 6. Briefly, siRNA and the DharmaFECT reagent were diluted in Opti-MEM, and the resulting siRNA-lipid complexes were added dropwise to the A549 cells in six-well plates seeded at 0.3 × 10^6^ cells per well to achieve a final siRNA concentration of 30 nM per well. The plates were gently shaken and incubated at 37 °C in a 5% CO_2_ incubator. After 8 h, the medium was replaced with a fresh complete medium. A non-targeting siRNA (siNT) was used as a negative control in all experiments. The cytotoxic effects associated with gene knockdown were evaluated at 72 h post-transfection (S5 Fig). NOXA knockdown was confirmed by western blot analysis of NOXA protein levels. IFNAR1 knockdown was inferred from reduced mRNA expression of *IFNAR1* and phosphorylation of STAT1.

**Table 6.**
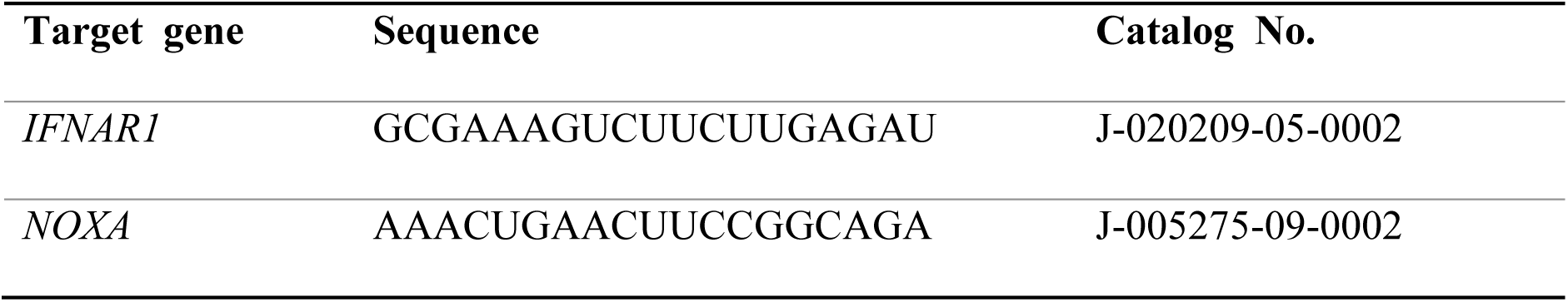
siRNA sequences used in this study.

### RNA-sequencing analysis

Total RNA was isolated from mock-treated (n = 2) and GAKV-infected A549 cells (n = 3) and submitted to Macrogen (Seoul, Republic of Korea) for library preparation and sequencing. Libraries were prepared using the TruSeq Stranded Total RNA Library Prep Gold Kit and sequenced in the paired-end mode on an Illumina NovaSeq platform. An average of 112 million paired-end reads were generated per sample. Raw sequencing reads were trimmed using Trimmomatic (v0.39) to remove adapter sequences and low-quality bases (sliding window: 4:15; leading/trailing: 3; minimum length: 36 bp). Trimmed reads were aligned to the human reference genome (GRCh38) using HISAT2. Gene-level expression counts were generated using the StringTie software for downstream analysis. The mapping rates ranged from 94%–98% across all samples. Differential expression analysis was performed using edgeR with TMM (trimmed mean of M value) normalization. Differentially expressed genes (DEGs) were defined as genes with |log_2_FC| ≥ 1 and Benjamini–Hochberg adjusted *P* < 0.05 compared to mock controls. Principal Component Analysis (PCA) and heat map visualization were performed using log_2_-transformed TMM-normalized values. Heatmaps were generated by row-wise Z-score normalization. Volcano plots were used to visualize the differential expression patterns, whereas violin plots were used to display the expression trends of representative antiviral and apoptosis-associated genes. Gene Ontology (GO) enrichment analysis was performed using the clusterProfiler R package (v4.6.2), focusing primarily on the Biological Process (BP) category, to investigate the biological pathways associated with GAKV infection. Significantly enriched GO biological processes were visualized as dot plots. GO terms with adjusted *P* < 0.05 were considered significantly enriched.

### Statistical analysis

All statistical analyses and graph generation were performed using GraphPad Prism, version 10.4.2. Data were analyzed for statistical significance using an unpaired two-tailed t-test, one-way ANOVA followed by Dunnett’s multiple-comparisons test, and two-way ANOVA followed by Tukey’s multiple-comparisons test as specified. All experiments were performed with three independent biological replicates. All statistical testing was performed at the two-sided alpha level of 0.05, and *P* < 0.05 was considered statistically significant.

## Results

### GAKV replication and innate immune activation in multiple cell lines

Multiple cell lines, including Vero E6, A549, and Huh7, were infected with GAKV to characterize viral replication and host responses. Samples were collected at 18, 24, 48, and 72 h post-infection (hpi) for analysis of cytotoxicity, viral replication, infectious virus production, and antiviral signaling (Fig 1A). Cytotoxicity analysis demonstrated a significant increase in cell death in A549 and Huh7 cells at 72 hpi, whereas minimal cytotoxicity was observed in Vero E6 cells (Fig 1B). In addition, syncytia were observed in GAKV-infected Vero E6 cells at 48 hpi, which increased by 72 hpi (S1 Fig). Quantification of intracellular viral RNA indicated active viral replication across all tested cell lines, with differences in replication kinetics (Fig 1C). Viral RNA levels progressively increased in Vero E6 and Huh7 cells, whereas A549 cells showed relatively stable viral RNA levels following the initial accumulation. Consistent with the viral RNA data, plaque assays demonstrated the production of infectious virus particles in all cell lines, with particularly high viral titers observed in Vero E6 and Huh7 cells (Fig 1D). Subsequently, *IFNB1* expression and activation of downstream signaling pathways were examined to characterize antiviral innate immune responses induced by GAKV infection. Robust induction of *IFNB1* expression was specifically observed in A549 cells, with peak expression detected at 24 hpi (Fig 1E). GAKV N protein was detected from 18 hpi, confirming infection (Fig 1F). Phosphorylation of IRF3 and STAT1 was observed with the strongest activation at 24-48 hpi. Consistent with the protein expression profile, *OASL* mRNA was also induced following GAKV infection (S2 Fig.). These findings indicated that A549 cells support GAKV replication while simultaneously mounting a measurable type I IFN response.

**Fig 1.**
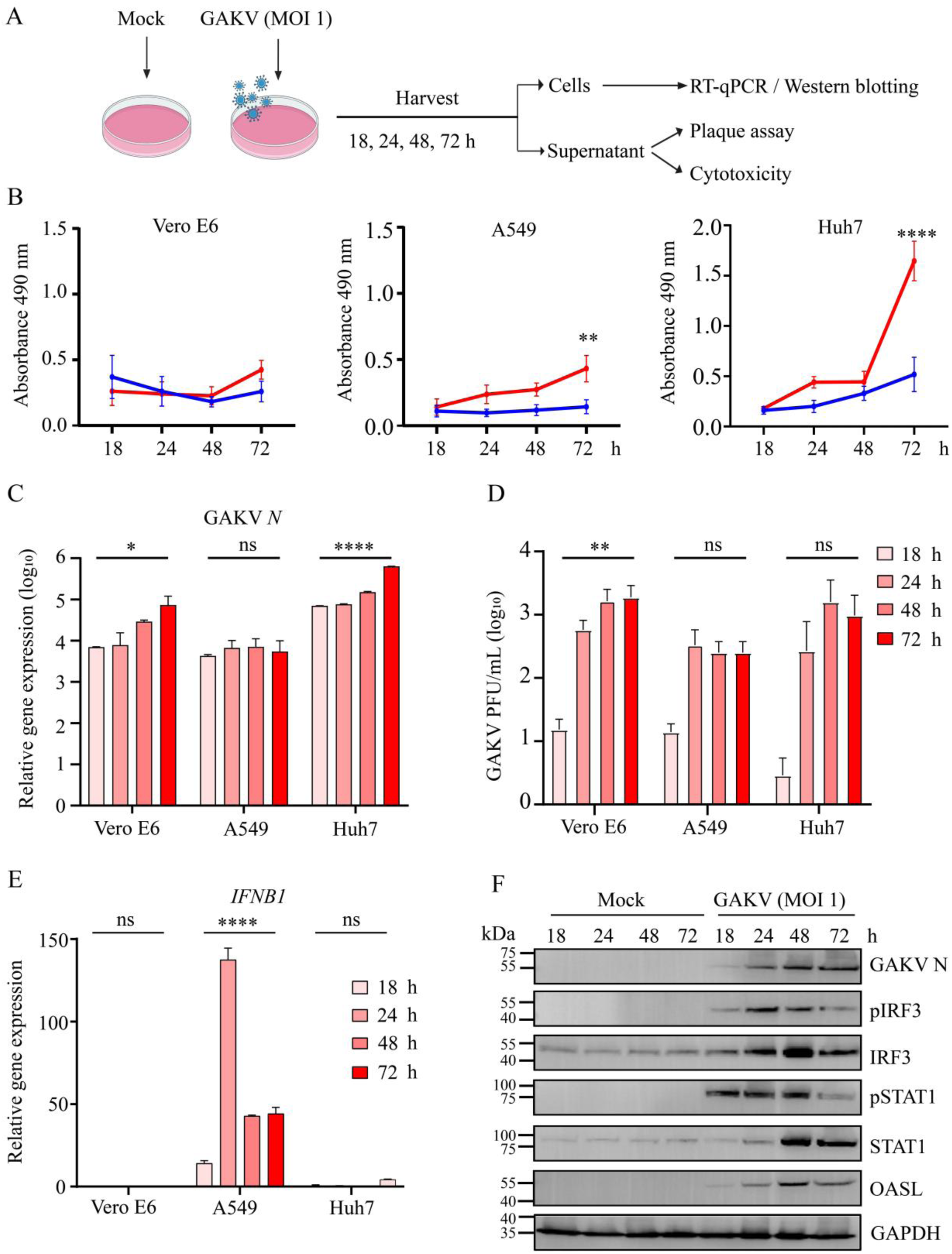
GAKV infects multiple cell lines but induces a robust innate immune response in A549 cells. **(A)** Experimental design showing Vero E6, A549, and Huh7 cells infected with GAKV (MOI 1) and harvested at 18, 24, 48, and 72 hours post-infection (hpi). Cell lysates were collected for reverse transcription-quantitative polymerase chain reaction (RT-qPCR) analysis, and A549 cell lysates were additionally analyzed by western blotting. Culture supernatants were used for plaque assays and cytotoxicity measurements. The image was created using bioRender. **(B)** Cell death was quantified by lactate dehydrogenase (LDH) release assay at the indicated time points. **(C)** GAKV *N* gene expression was quantified by RT-qPCR and normalized to GAPDH. **(D)** Viral titers in culture supernatants were determined by plaque assay and expressed as log_10_ PFU/mL. **(E)** Relative *IFNB1* mRNA expression was measured by RT-qPCR and normalized to GAPDH. **(F)** Immunoblot analysis of GAKV N, pIRF3, IRF3, pSTAT1, STAT1, and OASL proteins in mock and GAKV-infected A549 cells at the indicated time points. Data are presented as mean ± SD from three independent experiments. Statistical significance of LDH measurements in B was determined using two-way ANOVA with multiple-comparison testing. Statistical significance in C, D and E was determined by one-way ANOVA with Dunnett’s multiple-comparison test comparing infected cells at each time point with the corresponding 18 hpi group within each cell line. \**P* < 0.05, \*\**P* < 0.01, \*\*\**P* < 0.001, \*\*\*\**P* < 0.0001; ns, not significant.

Collectively, these results demonstrate that GAKV efficiently infects multiple cell lines and elicits distinct host responses. Among the evaluated cell lines, A549 cells supported viral replication and exhibited antiviral and cytotoxic responses. Therefore, A549 cells were selected for subsequent mechanistic studies to investigate the host response to GAKV infection.

### GAKV inoculation induces innate immune responses in BALB/c mice

Female BALB/c mice were intranasally inoculated with GAKV and euthanized at 1, 3, or 5 days post-infection (dpi) for analysis of lung viral RNA levels and host gene expression (Fig 2A). No significant differences in body weight were observed between the mock- and GAKV-inoculated animals throughout the experimental period, indicating the absence of overt clinical disease under the conditions tested (Fig 2B). GAKV *N* RNA was detected in lung tissues at 1 dpi and progressively declined at later time points (Fig 2C). Viral RNA was not detected in any other examined tissues (data not shown). The expression of representative antiviral and chemokine genes in the lungs was examined to assess innate immune responses following GAKV inoculation. Analysis of *Ifnb1* gene expression revealed an increase in GAKV-inoculated mice at 3 dpi, although the difference compared with time-matched mock controls did not reach statistical significance (Fig 2D). The ISGs *Ifit1* and *Isg15* were significantly upregulated at 1 dpi, with *Isg15* remaining significantly elevated at 3 dpi; expression of both genes declined by 5 dpi (Fig 2E, F). In addition, the expression of *Ccl5* and *Cxcl5* did not differ significantly between mock- and GAKV-inoculated animals at the examined time points (Fig 2G, H). Collectively, these findings demonstrate that intranasal GAKV inoculation results in transient detection of viral RNA in the lungs and the induction of antiviral host responses, despite the absence of clinical signs during the observation period.

**Fig 2.**
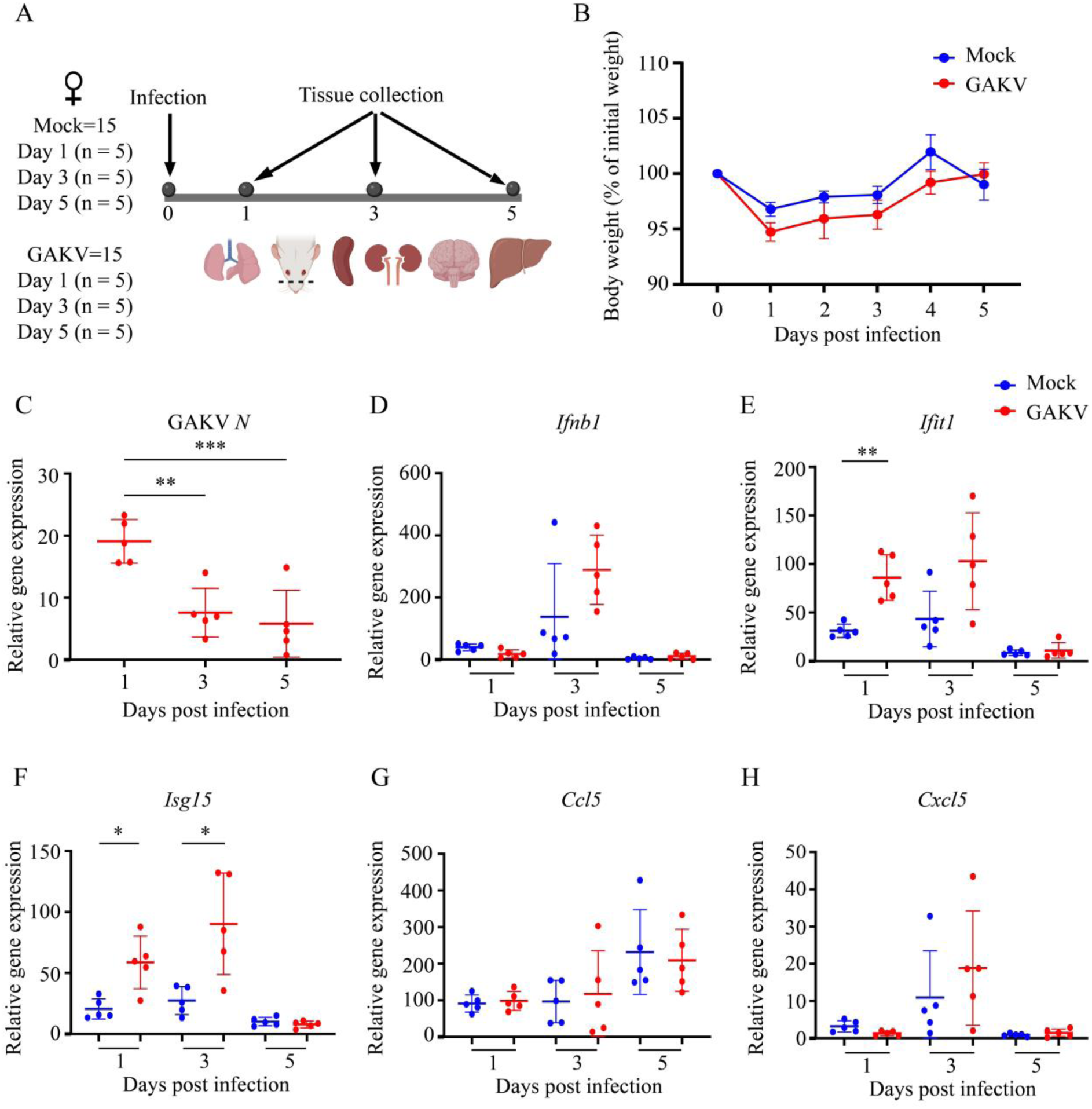
GAKV inoculation induces antiviral and inflammatory responses in the lungs of female BALB/c mice. **(A)** Experimental design showing intranasal inoculation of mice with GAKV and tissue collection at 1, 3, and 5 days post-infection (dpi). The image was created using bioRender. **(B)** Body weight changes following GAKV inoculation. Body weights are expressed as a percentage of the initial body weight. **(C)** Relative GAKV *N* RNA levels in lung tissues collected at the indicated time points. **(D-H)** Relative expression of *Ifnb1*, *Ifit1*, *Isg15*, *Ccl5*, and *Cxcl5* in lung tissues following GAKV inoculation. Gene expression was normalized to *Tuba1a*. Data are presented as mean ± SD. Differences between mock- and GAKV-inoculated mice at each time point were assessed using multiple unpaired two-tailed t-tests with Holm-Šídák correction for multiple comparisons. \**P* < 0.05, \*\**P* < 0.01, \*\*\**P* < 0.001, \*\*\*\**P* < 0.0001

### Transcriptomic profiling reveals activation of antiviral and apoptosis-associated gene programs during GAKV infection

Total RNA from GAKV-infected A549 cells at 48 hpi was subjected to RNA-seq-based host transcriptome analysis to determine the genome-wide gene expression profile. In infected samples, 2.9–3.6% of the total reads were mapped to the viral genome. Viral reads were distributed across all GAKV genes, including the nucleocapsid (*N*), phosphoprotein (*P*), matrix (*M*), fusion (*F*), receptor-binding protein (*RBP*), and large polymerase (*L*) genes. A transcriptional gradient was observed across the viral genome (S3 Fig). Principal component analysis (PCA) showed a clear separation between the mock- and GAKV-infected samples along PC1, which explained 96.52% of the total variance (Fig 3A). Differential expression analysis identified 2,064 genes significantly altered upon infection, including 1,454 upregulated and 610 downregulated genes, indicating a robust host transcriptional response. Furthermore, Gene Ontology (GO) enrichment analysis revealed strong activation of antiviral and innate immune pathways, with “response to virus” representing the most significantly enriched biological process (Fig 3B). Additional enriched pathways included positive regulation of cytokine production, NF-κB signaling, cellular response to type I IFN, and intrinsic apoptotic signaling, indicating that GAKV infection induces coordinated antiviral and cell death-associated transcriptional programs. Consistent with these findings, volcano plot analysis demonstrated robust upregulation of multiple antiviral ISGs, including *MX1*, *OAS1*, *OAS2*, *OASL*, *IFIT1*, and *IFIT2,* together with the interferon-responsive gene *XAF1*. Additionally, apoptosis-associated genes, including *NOXA* and *PUMA*, were induced following infection (Fig 3C). Heatmap visualization and normalized transcript quantification further demonstrated clear segregation between mock- and GAKV-infected samples based on the expression of representative ISGs and apoptosis-related genes (Fig 3D, E). Collectively, these findings indicate that GAKV infection induces a dominant type I IFN-associated antiviral transcriptional response, accompanied by the activation of genes associated with intrinsic apoptosis, suggesting coordinated induction of antiviral and apoptotic programs during infection.

**Fig 3.**
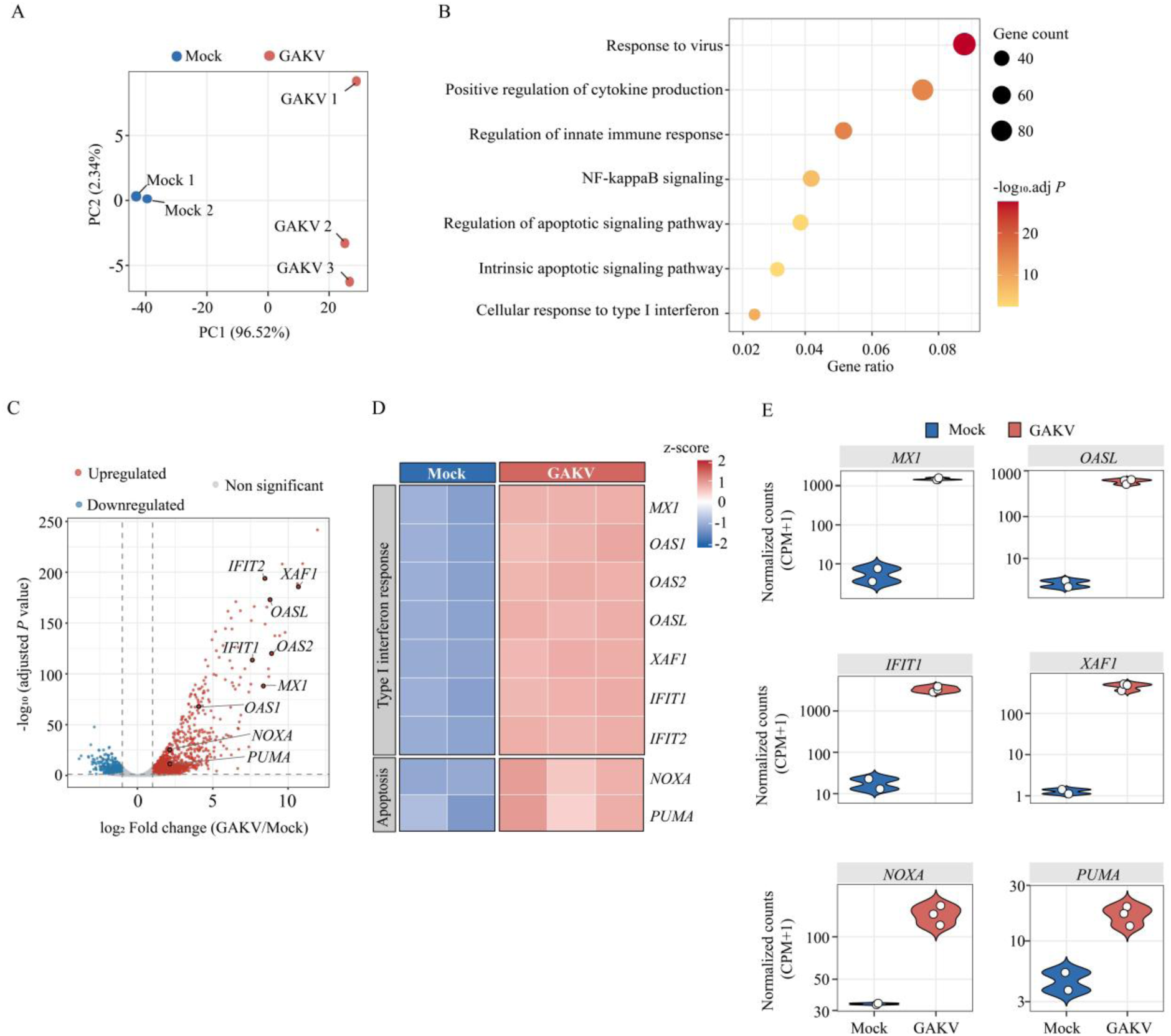
GAKV infection induces antiviral and pro-apoptotic transcriptional responses in A549 cells. RNA-seq analysis was performed using total RNA isolated from mock-treated (n = 2) and GAKV-infected A549 cells (n = 3) at 48 hours post-infection (hpi). **(A)** Principal component analysis (PCA) of global gene expression profiles showing mock and GAKV-infected samples based on trimmed mean of M value (TMM)-normalized counts per million (CPM) values. Each point represents an individual sample, and the percentage of variance explained by each principal component is indicated on the axes. **(B)** Gene Ontology (GO) enrichment analysis of upregulated genes revealed significant enrichment of biological processes in GAKV-infected A549 cells at 48 hpi compared with mock-infected controls. Dot size represents the number of genes associated with each pathway, while dot color indicates enrichment significance (-log_10_ adjusted *P* value). **(C)** Volcano plot showing differentially expressed genes (DEGs) in GAKV-infected A549 cells compared to mock-infected controls. Each dot represents an individual gene. Red dots indicate significantly upregulated genes, blue dots indicate significantly downregulated genes, and gray dots indicate non-significant genes. Representative interferon-stimulated genes (ISGs) and apoptosis-associated genes are highlighted. **(D)** Heatmap showing expression patterns of type I interferon-responsive and apoptosis-associated genes in mock- and GAKV-infected samples. Expression values are displayed as row-wise Z-scores of TMM-normalized transcript abundance. **(E)** Violin plots showing TMM-normalized transcript abundance of representative type I interferon-responsive and apoptosis-associated genes in mock- and GAKV-infected samples. DEGs were identified using edgeR with Benjamini–Hochberg adjusted *P* value < 0.05 and |log_2_ FC| > 1.

### GAKV infection induces apoptosis in A549 cells

The activation of apoptotic markers was examined at multiple time points following infection to determine whether GAKV infection induces apoptosis. Western blot analysis revealed progressive activation of apoptotic signaling during GAKV infection (Fig 4A). Cleavage of caspase-8 and caspase-9 was detected at 24 hpi and increased at 72 hpi, whereas cleaved caspase-3 was observed from 48 hpi. These findings demonstrated activation of both initiator and executioner caspases during GAKV infection. Subsequently, cleaved caspase-3 expression was examined using immunofluorescence microscopy to further assess apoptosis at the single-cell level (Fig 4B). Quantification of cleaved caspase-3 fluorescence intensity revealed a significant increase at 48 hpi, followed by a further increase at 72 hpi (Fig 4C). Phosphatidylserine (PS) exposure was examined using Apopxin Green staining and combined with GAKV N protein detection to determine whether apoptotic cells corresponded to virus-infected cells (Fig 4D). Apopxin-positive cells were detected in the GAKV-infected cultures at 48 hpi, whereas staining was absent in the mock-infected cells. Co-staining for the GAKV N protein showed overlap between Apopxin-positive and virus-infected cells, indicating that apoptotic cell death occurred predominantly within infected cell populations. Collectively, these findings demonstrate that GAKV infection is associated with progressive activation of apoptotic markers, including caspase cleavage and PS externalization.

**Fig 4.**
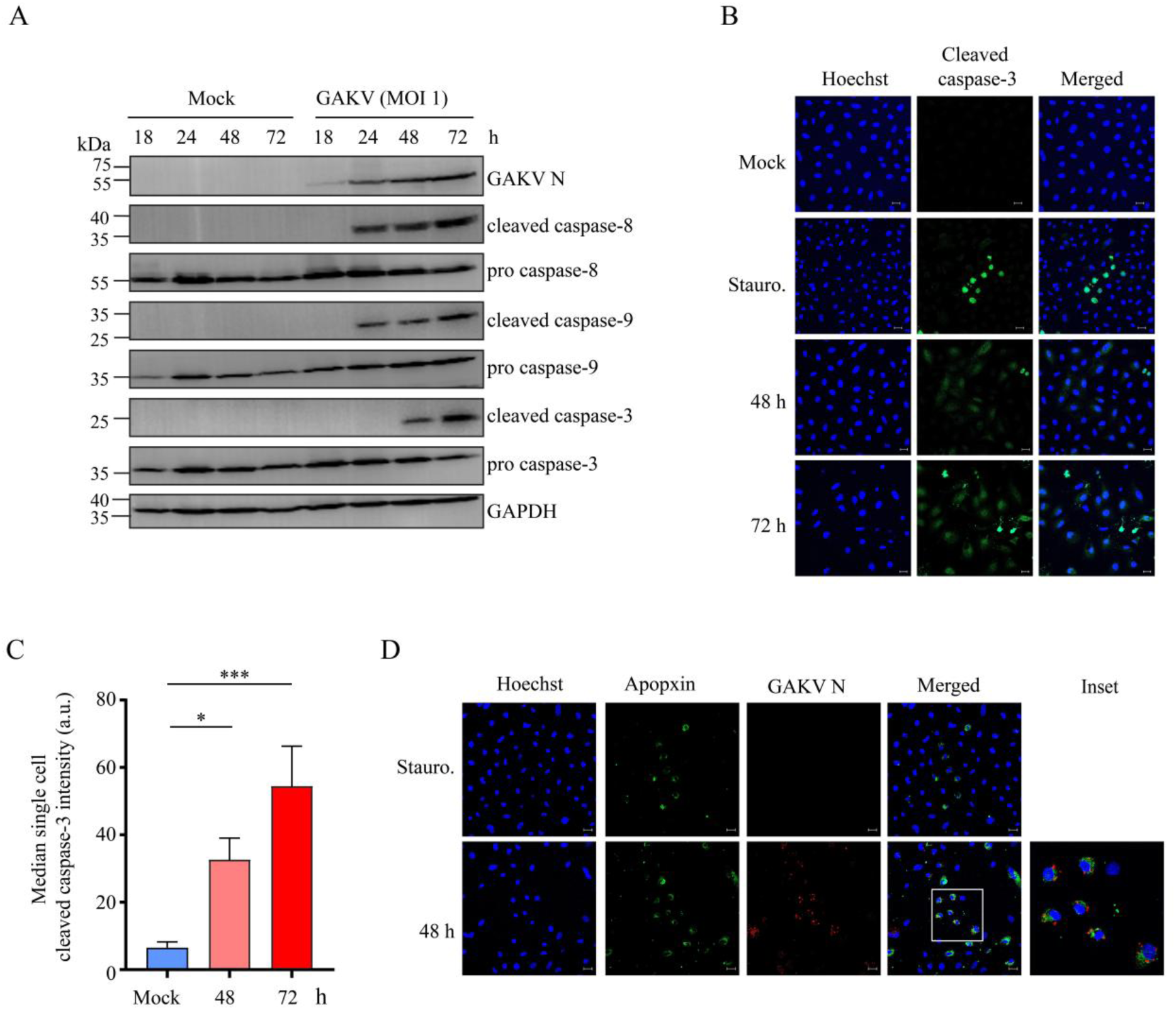
Induction of apoptosis during GAKV infection. **(A)** A549 cells were infected with GAKV (MOI 1) and harvested at the indicated time points post-infection. GAKV nucleocapsid (N) protein expression and cleavage of caspase-8, caspase-9, and caspase-3 were assessed by western blotting. Corresponding procaspase proteins are shown for comparison. GAPDH was used as a loading control. **(B)** A549 cells were stained for cleaved caspase-3 (green) and nuclei (Hoechst, blue). Representative images of mock-treated cells, staurosporine-treated cells (Stauro., positive control), and GAKV-infected cells at 48 and 72 hpi are shown. Scale bars, 20 μm. **(C)** Cleaved caspase-3 fluorescence intensity was measured and is presented as the median single-cell fluorescence intensity in arbitrary units (a.u.). Data represent mean ± SD from three independent experiments. Statistical significance was determined using one-way analysis of variance (ANOVA) followed by Dunnett’s multiple comparisons test using mock-treated cells as the reference group. **(D)** Detection of Phosphatidylserine (PS) exposure during GAKV infection using Apopxin Green staining. A549 cells were stained with Apopxin Green (green), Hoechst (blue), and an antibody against the GAKV N protein (red) at 48 hpi. The inset shows a magnified view of the indicated region. Scale bars, 20 μm.

### RIG-I–MAVS signaling restricts GAKV replication and promotes apoptosis during infection

A549-derived RIG-I knockout (KO), MDA5 KO, and MAVS KO cells were infected with GAKV and analyzed for viral replication, antiviral signaling, and apoptosis to investigate the role of RLR signaling during infection. Immunofluorescence staining of the GAKV N protein revealed an increased proportion of viral antigen-positive cells in RIG-I KO and MAVS KO cultures compared with gNT control cells, whereas MDA5 KO cells exhibited infection levels comparable to those of the controls (Fig 5A). Quantification of GAKV N-positive cells confirmed significantly enhanced infection in RIG-I KO and MAVS KO cells, whereas no significant difference was observed in MDA5 KO cells (Fig 5B). Consistent with these findings, plaque assays showed a significantly increased production of infectious virus particles in RIG-I KO and MAVS KO cells compared to gNT cells, whereas viral titers in MDA5 KO cells remained comparable to those in gNT cells (Fig 5C). These results indicate that RIG-I–MAVS signaling plays a dominant role in restricting GAKV replication. Subsequently, we assessed the activation of downstream signaling molecules by western blot analysis to determine whether antiviral signaling was dependent on RLR pathways. GAKV infection induced phosphorylation of IRF3, IκBα, STAT1, and expression of the ISG OASL in gNT and MDA5 KO cells (Fig 5D). In contrast, activation of these signaling pathways was markedly absent in RIG-I KO and MAVS KO cells, indicating that antiviral signaling induced by GAKV is primarily mediated through the RIG-I–MAVS axis.

**Fig 5.**
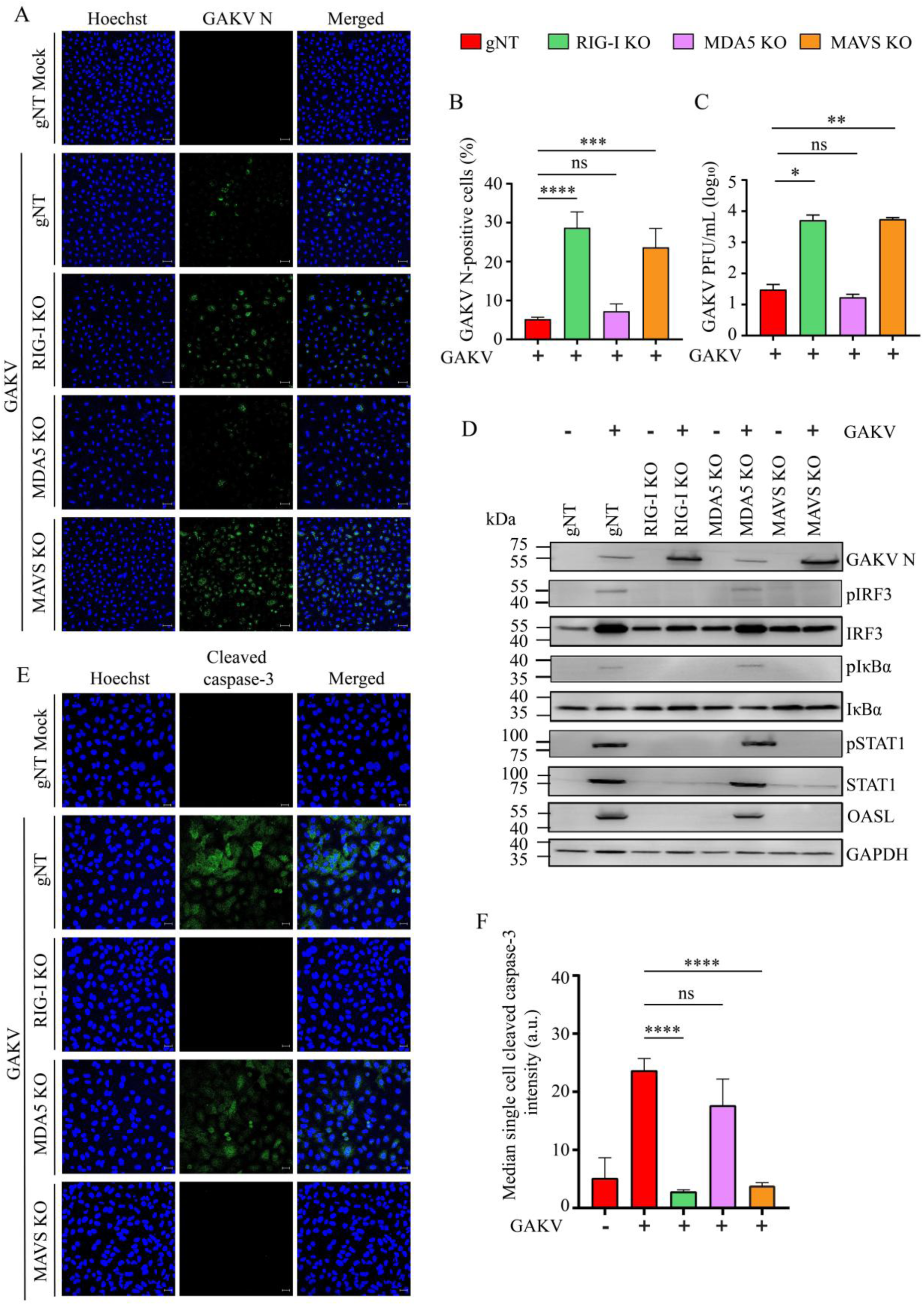
RIG-I-MAVS signaling regulates GAKV infection, antiviral signaling, and apoptosis in A549 cells. **(A)** Immunofluorescence analysis of GAKV infection in gNT, RIG-I knockout (KO), MDA5 KO, and MAVS KO A549 cells at 48 hours post-infection (hpi). Cells were stained for GAKV N protein (green), and nuclei were counterstained with Hoechst (blue). Scale bars, 10 µm. **(B)** Quantification of GAKV N-positive cells expressed as the percentage of total cells. **(C)** Viral titers in culture supernatants at 48 hpi were determined by plaque assay and expressed as log_10_ PFU/mL. **(D)** Western blot analysis of GAKV N, pIRF3, IRF3, pIκBα, IκBα, pSTAT1, STAT1, and OASL expression in gNT, RIG-I KO, MDA5 KO, and MAVS KO cells following GAKV infection at 48 hpi. GAPDH was used as a loading control. **(E)** Immunofluorescence staining of cleaved caspase-3 in infected gNT, RIG-I KO, MDA5 KO, and MAVS KO cells at 48 hpi. Cells were stained for cleaved caspase-3 (green), and nuclei were counterstained with Hoechst (blue). Scale bars, 20 µm. **(F)** Quantification of the cleaved caspase-3 signal at the single-cell level is represented as median fluorescence intensity (a.u.). Data are presented as mean ± SD from three independent experiments. Statistical significance was determined using one-way analysis of variance (ANOVA) followed by Dunnett’s multiple-comparison test. \**P* < 0.05, \*\**P* < 0.01, \*\*\**P* < 0.001, \*\*\*\**P* < 0.0001; ns, not significant.

Thereafter, we examined whether RLR signaling contributed to apoptotic responses. Immunofluorescent staining for cleaved caspase-3 demonstrated robust apoptosis in GAKV-infected gNT cells (Fig 5E). In contrast, disruption of RIG-I or MAVS markedly reduced cleaved caspase-3 staining, whereas MDA5 KO cells exhibited levels comparable to those in gNT cells. Quantification of single-cell cleaved caspase-3 fluorescence intensity confirmed a significant reduction in apoptosis in RIG-I KO and MAVS KO cells relative to that in infected gNT cells (Fig 5F). Collectively, these findings identify RIG-I as the primary sensor of GAKV infection in A549 cells and demonstrate that RIG-I–MAVS signaling is required for antiviral innate immune activation and the induction of apoptosis during infection.

### Type I IFN signaling contributes to antiviral defense and apoptosis during GAKV infection

*IFNAR1* expression was reduced by siRNA-mediated knockdown in A549 cells to investigate whether type I interferon (IFN) signaling contributes to apoptotic responses during GAKV infection. RT-qPCR analysis confirmed the efficient knockdown of *IFNAR1* expression in siIFNAR1-treated cells compared to that in siNT-treated cells (Fig 6A). Consistent with impaired type I IFN signaling, induction of the ISG *OAS1* was significantly reduced following GAKV infection in siIFNAR1-treated cells (Fig 6B). This was accompanied by a significant increase in infectious virus production (Fig 6C). Western blot analysis further demonstrated reduced STAT1 phosphorylation and increased accumulation of GAKV N protein in siIFNAR1-treated cells compared with siNT controls. In parallel, cleaved caspase-3 expression was markedly reduced following IFNAR1 knockdown, indicating that disruption of type I IFN signaling attenuates apoptosis during GAKV infection (Fig 6D).

**Fig 6.**
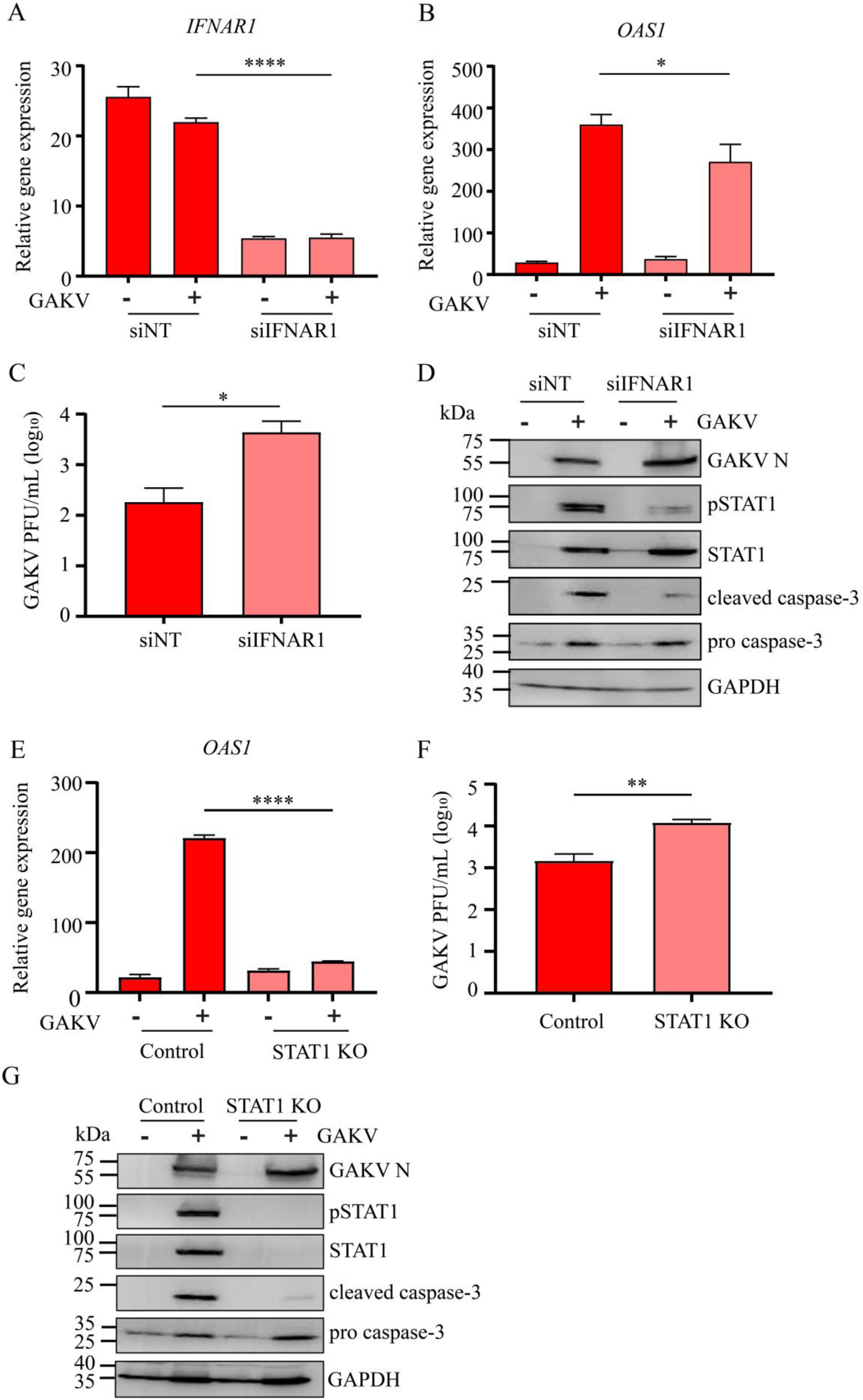
Disruption of type I IFN signaling enhances GAKV replication and attenuates apoptosis during GAKV infection. (A–D) A549 cells were transfected with non-targeting control siRNA (siNT) or IFNAR1-targeting siRNA (siIFNAR1) and infected with GAKV at an MOI of 1. **(A)** Relative *IFNAR1* mRNA expression normalized to GAPDH. **(B)** Relative *OAS1* mRNA expression following GAKV infection. **(C)** Viral titers in culture supernatants at 48 h post-infection (hpi), expressed as log_10_ PFU/mL. **(D)** Western blot analysis of GAKV N, pSTAT1, STAT1, cleaved caspase-3 and pro caspase-3 expression. GAPDH was used as a loading control. **(E–G)** Control and STAT1 knockout (KO) A549 cells were infected with GAKV at an MOI of 1. **(E)** Relative *OAS1* mRNA expression normalized to GAPDH. **(F)** Viral titers in culture supernatants at 48 hpi, expressed as log_10_ PFU/mL. **(G)** Western blot analysis of GAKV N, pSTAT1, STAT1, cleaved caspase-3 and pro caspase-3 expression. GAPDH was used as a loading control. Data are presented as mean ± SD from three independent experiments. Statistical significance was determined using an unpaired two-tailed Student’s t-test. \**P* < 0.05, \*\**P* < 0.01, \*\*\**P* < 0.001, \*\*\*\**P* < 0.0001.

To further validate the role of downstream IFN signaling, STAT1 KO cells were examined following GAKV infection. Similar to IFNAR1 knockdown, STAT1 deficiency markedly reduced *OAS1* induction (Fig 6E) and significantly increased infectious virus production (Fig 6F). Western blot analysis confirmed the absence of STAT1 expression and demonstrated reduced cleaved caspase-3 levels despite productive infection, as indicated by GAKV N protein expression (Fig 6G). Collectively, these findings demonstrate that type I IFN signaling promotes both antiviral defense and apoptosis during GAKV infection. IFNAR1 knockdown and STAT1 deficiency provide complementary evidence that disruption of the type I IFN signaling pathway enhances viral replication while suppressing apoptotic responses.

### NOXA contributes to GAKV-induced apoptosis downstream of innate immune signaling

We examined NOXA expression to further investigate apoptotic signaling during GAKV infection. RT-qPCR analysis demonstrated the progressive induction of *NOXA* expression following infection, with transcript levels increasing throughout the infection period (Fig 7A). Subsequently, *NOXA* expression was examined in RLR-deficient cells to determine whether its induction occurs downstream of RLR signaling during GAKV infection. GAKV infection induced robust *NOXA* expression in gNT and MDA5 KO cells; however, this induction was significantly reduced in RIG-I KO and MAVS KO cells (Fig 7B). These findings indicate that the upregulation of *NOXA* during GAKV infection requires the RIG-I-MAVS signaling axis. GAKV-induced *NOXA* expression was significantly reduced in both siIFNAR1-treated and STAT1 KO cells (Fig. 7C, D), indicating that NOXA induction depends on intact type I IFN signaling downstream of viral sensing.

**Fig 7.**
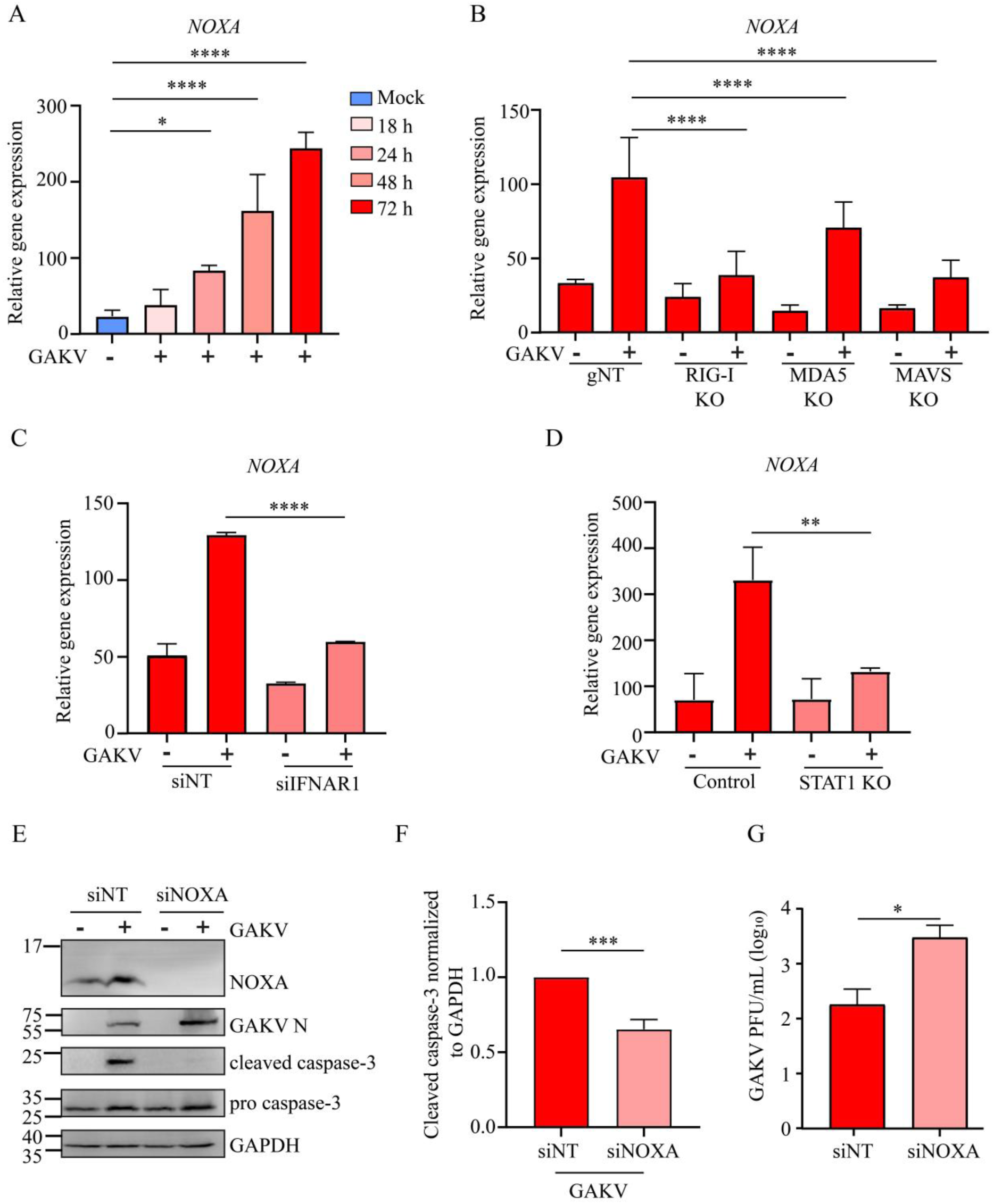
NOXA contributes to apoptotic responses during GAKV infection. **(A)** Relative *NOXA* mRNA expression in GAKV-infected A549 cells at 18, 24, 48, and 72 hours post-infection (hpi). **(B)** Relative *NOXA* mRNA expression in GAKV-infected gNT, RIG-I KO, MDA5 KO, and MAVS KO A549 cells at 48 hpi. **(C)** Relative *NOXA* mRNA expression in siNT- and siIFNAR1-transfected A549 cells following GAKV infection. **(D)** Relative *NOXA* mRNA expression in CRISPR Control (Control) and STAT1 KO A549 cells following GAKV infection. Gene expression for RT-qPCR was normalized to GAPDH. **(E)** Western blot analysis of NOXA, pro caspase-3, cleaved caspase-3, and GAKV N protein expression in siNT- and siNOXA-transfected A549 cells following GAKV infection. GAPDH was used as a loading control. **(F)** Quantification of cleaved caspase-3 protein levels normalized to GAPDH from western blot analysis. **(G)** Viral titers in culture supernatants were determined by plaque assay and expressed as log_10_ PFU/mL. Data are presented as mean ± SD from three independent experiments. Statistical significance was determined using one-way analysis of variance (ANOVA) followed by Dunnett’s multiple-comparison test for panels A and B and an unpaired two-tailed Student’s t-test for panels C, D, F and G. \**P* < 0.05, \*\**P* < 0.01, \*\*\**P* < 0.001, \*\*\*\**P* < 0.0001.

To investigate whether NOXA functionally contributes to apoptosis during GAKV infection, A549 cells were transfected with either siNT or siNOXA prior to infection. Western blot analysis confirmed efficient depletion of NOXA protein in siNOXA-transfected cells and demonstrated a marked reduction in cleaved caspase-3 levels compared with infected siNT controls (Fig. 7E). Quantification of cleaved caspase-3 normalized to GAPDH confirmed a significant reduction in caspase-3 activation following NOXA knockdown (Fig. 7F), indicating that NOXA promotes apoptotic signaling during GAKV infection. In addition, NOXA depletion increased GAKV N protein expression and significantly increased infectious virus production compared with siNT-transfected cells (Fig. 7E, G). Collectively, these findings identify NOXA as a downstream effector of innate immune signaling that promotes apoptosis and contributes to restricting GAKV replication.

## Discussion

The emergence of LayV in humans highlights the importance of defining host responses to infection by members of the genus *Parahenipavirus*. GAKV induces antiviral responses in A549 cells during the early stages of infection; the pathways responsible for viral sensing and the biological consequences of antiviral signaling remain unclear [7]. In this study, we identified RIG-I as the primary sensor of GAKV infection and that RIG-I–MAVS-dependent signaling contributes to apoptotic responses through the downstream induction of NOXA. Together, these findings provide the first mechanistic characterization of innate immune sensing and apoptosis induced by a parahenipavirus, establishing GAKV as a tractable model for investigating host responses to this emerging genus.

GAKV infection elicited a robust antiviral response in A549 cells, characterized by IRF3 and STAT1 activation, increased *IFNB1* and OASL expression, indicating efficient engagement of innate immune sensing pathways. Similar induction of type I IFN has been reported during NiV infection in primary human endothelial cells [22]. However, unlike NiV, which continues to replicate efficiently despite activation of innate immune responses because of its potent immune evasion strategies. The antiviral response induced by GAKV was associated with more restricted viral replication in A549 cells than in Huh7 cells, suggesting that efficient innate immune activation contributes to limiting GAKV replication in IFN-competent cells. Although multiple host cell factors likely contribute to these differences, the enhanced viral replication observed following disruption of type I IFN signaling supports the role of innate immune responses in limiting GAKV infection.

GAKV exposure in wild-type BALB/c mice elicited antiviral responses in the absence of overt clinical symptoms. Although productive GAKV replication was not observed, detection of viral RNA in the lungs, together with the induction of antiviral innate immune genes, indicates that intranasal GAKV exposure is sufficient to activate host antiviral recognition pathways *in vivo*. A similar outcome has been reported following NiV infection in BALB/c mice, in which infection remained largely restricted to the respiratory tract and was rapidly controlled by host antiviral responses without extensive systemic dissemination or neurological diseases [23]. Chemokines contribute to antiviral immunity by coordinating the recruitment of immune cells to sites of infection [24]. GAKV inoculation did not significantly alter the expression of *Ccl5* or *Cxcl5,* which are associated with the recruitment of monocytes/lymphocytes and neutrophils, respectively [25, 26]. Collectively, these findings support a model in which innate immune activation contributes to the early control of GAKV infection in mice without pronounced changes in the inflammatory chemokines examined in the lung.

Transcriptomic analysis revealed that GAKV infection elicits a coordinated host response involving antiviral and apoptotic signaling pathways. Although the enrichment of type I IFN pathways and ISG expression was consistent with the observed antiviral response, the simultaneous enrichment of intrinsic apoptotic pathways suggests that cell death forms an integrated component of the host response to infection. Notably, *NOXA* and *PUMA*, two BH3-only regulators of intrinsic apoptosis, are among the genes induced following infection [27]. The induction of these genes suggests that antiviral signaling and apoptotic responses may be coordinately regulated during GAKV infection. Subsequent genetic and functional analyses supported this hypothesis and identified a signaling pathway linking antiviral sensing to apoptotic signaling.

RIG-I has been shown to mediate innate immune responses during paramyxoviral infections [28, 29]. Furthermore, the requirement for MAVS observed in this study is consistent with previous evidence implicating MAVS in restricting NiV infections *in vitro* and *in vivo* [13]. Our findings further extend the role of the RIG-I–MAVS signaling axis beyond antiviral defense by demonstrating its contribution to apoptotic responses during GAKV infection. The requirement for RIG-I and MAVS, but not MDA5, indicates that RIG-I–MAVS signaling serves as the primary viral-sensing pathway linking GAKV detection to apoptotic responses. Notably, the disruption of either RIG-I or MAVS reduced antiviral signaling and caspase-3 activation, supporting a functional connection between innate immune activation and apoptotic responses. Similar links between RIG-I signaling and apoptosis have been described in multiple viral systems. For instance, during Sendai virus (SeV) infection, RIG-I can promote cell death through the RIG-I-induced IRF3-mediated apoptosis pathway (RIPA), a mechanism that occurs independently of NF-κB or IRF3 transcriptional activation [30]. In addition, RIG-I-dependent signaling induces the expression of the BH3-only protein, NOXA, during reovirus infection, linking innate immune sensing to intrinsic apoptotic signaling [31]. MAVS has also been implicated in regulating apoptosis during infection with SeV, dengue virus, and Semliki Forest virus and has been proposed to coordinate antiviral apoptotic responses at the mitochondrial membrane [32–34]. Consistent with the broad relevance of apoptosis during paramyxoviral infection, caspase-3 activation has been reported in HeV-infected A549 cells [2].

Type I IFN signaling contributes to viral restriction and apoptosis during GAKV infection. Both IFNAR1 knockdown and STAT1 KO attenuated apoptotic signaling, supporting an important role for type I IFN-STAT1 signaling in promoting apoptosis during GAKV infection. However, residual apoptotic signaling remained detectable in STAT1-deficient cells, indicating that additional IFN-independent pathways may also contribute to apoptosis [35]. Consistent with previous reports showing that IFN signaling induces pro-apoptotic gene expression and sensitizes cells to apoptosis, we sought to identify the downstream effectors linking antiviral signaling to apoptosis [36, 37]. NOXA has previously been reported to be induced during infection with multiple RNA viruses and contribute to antiviral apoptotic responses [38, 39]. Consistent with these observations, GAKV infection induced *NOXA* expression, whereas the disruption of RIG-I, MAVS, IFNAR1 or STAT1 substantially reduced *NOXA* induction. Notably, NOXA knockdown enhanced viral replication, suggesting that NOXA-dependent apoptotic signaling contributes to the restriction of GAKV replication and spread. Together, these findings support a model in which RIG-I-MAVS-dependent sensing initiates antiviral signaling, type I IFN-STAT1 axis promotes NOXA induction, and NOXA functions as a downstream effector linking innate immune activation to the apoptotic elimination of infected cells.

However, this study has some limitations that should be considered when interpreting the findings. Although our data support a role for apoptosis in GAKV infection, the relative contributions of intrinsic and extrinsic apoptotic pathways remain unclear. Cleavage of both caspase-8 and caspase-9 was observed during infection, indicating the activation of multiple caspase-dependent pathways. While NOXA induction and caspase-9 activation support the involvement of the intrinsic pathway, cleavage of caspase-8 may reflect either direct activation of extrinsic apoptotic signaling or secondary amplification downstream of executioner caspases. In addition, the molecular mechanisms linking NOXA induction to mitochondrial dysfunction were not directly examined. Future studies assessing mitochondrial membrane potential, cytochrome c release, and other downstream mitochondrial events will be required to define further the apoptotic pathway activated during GAKV infection. Although PUMA was also induced during GAKV infection, its functional contribution was not examined in the present study. PUMA may act independently of or in concert with NOXA to promote mitochondrial apoptosis, and future studies will be required to determine whether it also contributes to GAKV-induced cell death and to the restriction of viral replication. In the mouse studies, histopathological analysis was not performed; therefore, the relationship between immune gene expression and tissue-level pathology could not be directly evaluated. Furthermore, viral dissemination and clearance were assessed primarily by quantifying viral RNA, which does not distinguish between productive replication and the persistence of residual viral genomes. Although GAKV exposure induces antiviral responses, the cellular sources of responses and the extent of immune cell recruitment remain uncharacterized. Further studies in additional animal models and host systems are needed to understand the *in vivo* pathogenesis and host responses to GAKV infection. Nevertheless, our findings establish a mechanistic link between innate immune sensing and apoptosis during GAKV infection, and provide a foundation for future studies investigating viral pathogenesis, host adaptation, and disease outcomes associated with emerging parahenipaviruses.

## Acknowledgments

We thank Nayeon Jang (Hallym University) for support in this study. We also thank Dr. Susan Weiss for providing the CRISPR control and STAT1 knockout (KO) A549 cells.

## Author Contributions Statement

**Shivani Rajoriya**: Conceptualization, Data curation, Investigation, Methodology, Writing – original draft. **Divya Misra**: Data curation, Methodology, Investigation. **Sun Hye Yu**: Software, Data curation, Investigation. **Altanzul Bat Ulzii**: Investigation, Visualization. **Hennisa Hennisa**: Investigation, Visualization. **Tae-Wook Kang**: Resources, Software. **Hye Jin Shin**: Validation, Visualization. **Yeonsu Oh**: Resources, Visualization. **Carolina Lopez**: Analysis, review and funding acquisition. **Won-Keun Kim**: Writing – review & editing, Project administration, Funding acquisition, Supervision

## Data Availability Statement

The source data supporting the findings of this study are openly available in Figshare at DOI: https://doi.org/10.6084/m9.figshare.33193935.

The RNA-seq data generated in this study have been deposited in the NCBI Sequence Read Archive (SRA) under BioProject accession number PRJNA1503114 and are publicly available at https://www.ncbi.nlm.nih.gov/bioproject/PRJNA1503114.

## References

1. Duprex, W.P. and R.E. Dutch, Paramyxoviruses: Pathogenesis, Vaccines, Antivirals, and Prototypes for Pandemic Preparedness. The Journal of Infectious Diseases, 2023. 228(Supplement_6): p. S390-S397.

2. Wynne, J.W., et al., Proteomics informed by transcriptomics reveals Hendra virus sensitizes bat cells to TRAIL-mediated apoptosis. Genome Biology, 2014. 15(11): p. 532.

3. Noh, J.Y., et al., Isolation and characterization of novel bat paramyxovirus B16-40 potentially belonging to the proposed genus Shaanvirus. Scientific Reports, 2018. 8(1): p. 12533.

4. de Souza, W.M., et al., Paramyxoviruses from neotropical bats suggest a novel genus and nephrotropism. Infection, Genetics and Evolution, 2021. 95: p. 105041.

5. Park, J.Y., et al., Detection, isolation, and in vitro characterization of porcine parainfluenza virus type 1 isolated from respiratory diagnostic specimens in swine. Veterinary Microbiology, 2019. 228: p. 219–225.

6. Natasha, A., et al., Detection and characterization of Langya virus in Crocidura lasiura (the Ussuri white-toothed shrew), Republic of Korea. One Health, 2025. 20: p. 101017.

7. Lee, S.-H., et al., Discovery and Genetic Characterization of Novel Paramyxoviruses Related to the Genus Henipavirus in Crocidura Species in the Republic of Korea. Viruses, 2021. 13(10): p. 2020.

8. Zhang, X.-A., et al., A Zoonotic Henipavirus in Febrile Patients in China. New England Journal of Medicine, 2022. 387(5): p. 470–472.

9. Kumagai, Y. and S. Akira, Identification and functions of pattern-recognition receptors. Journal of Allergy and Clinical Immunology, 2010. 125(5): p. 985–992.

10. Rehwinkel, J. and M.U. Gack, RIG-I-like receptors: their regulation and roles in RNA sensing. Nature Reviews Immunology, 2020. 20(9): p. 537–551.

11. Yoneyama, M., H. Kato, and T. Fujita, Physiological functions of RIG-I-like receptors. Immunity, 2024. 57(4): p. 731–751.

12. Melchjorsen, J., et al., Activation of Innate Defense against a Paramyxovirus Is Mediated by RIG-I and TLR7 and TLR8 in a Cell-Type-Specific Manner. Journal of Virology, 2005. 79(20): p. 12944–12951.

13. Iampietro, M., et al., Control of Nipah Virus Infection in Mice by the Host Adaptors Mitochondrial Antiviral Signaling Protein (MAVS) and Myeloid Differentiation Primary Response 88 (MyD88). The Journal of Infectious Diseases, 2020. 221(Supplement_4): p. S401-S406.

14. Melchjorsen, J., L.N. Sørensen, and S.R. Paludan, Expression and function of chemokines during viral infections: from molecular mechanisms to in vivo function. Journal of Leukocyte Biology, 2003. 74(3): p. 331–343.

15. Riera Romo, M., Cell death as part of innate immunity: Cause or consequence? Immunology, 2021. 163(4): p. 399–415.

16. Everett, H. and G. McFadden, Apoptosis: an innate immune response to virus infection. Trends in Microbiology, 1999. 7(4): p. 160–165.

17. Danthi, P., Viruses and the Diversity of Cell Death. Annual Review of Virology, 2016. 3(Volume 3, 2016): p. 533-553.

18. Gupta, M., M.K. Lo, and C.F. Spiropoulou, Activation and cell death in human dendritic cells infected with Nipah virus. Virology, 2013. 441 **1**: p. 49–56.

19. Lin, Y., et al., Induction of apoptosis by paramyxovirus simian virus 5 lacking a small hydrophobic gene. Journal of Virology, 2003. 77(6): p. 3371–83.

20. Esolen, L.M., et al., Apoptosis as a cause of death in measles virus-infected cells. Journal of Virology, 1995. 69(6): p. 3955–3958.

21. Whelan, J.N., et al., Zika Virus Production Is Resistant to RNase L Antiviral Activity. Journal of Virology, 2019. 93(16).

22. Lo, M.K., et al., Characterization of the antiviral and inflammatory responses against Nipah virus in endothelial cells and neurons. Virology, 2010. 404(1): p. 78–88.

23. Dups, J., et al., Subclinical infection without encephalitis in mice following intranasal exposure to Nipah virus-Malaysia and Nipah virus-Bangladesh. Virology Journal, 2014. 11(1): p. 102.

24. Pita-Martínez, C., et al., CCL5 Levels, Disease Progression, and Mortality in Respiratory Viral Infections: A Systematic Review and Meta-Analysis. JAMA Network Open, 2026. 9(8): p. e2628987–e2628987.

25. Culley, F.J., et al., Role of CCL5 (RANTES) in viral lung disease. Journal of Virology, 2006. 80(16): p. 8151–7.

26. Guo, L., et al., Role of CXCL5 in Regulating Chemotaxis of Innate and Adaptive Leukocytes in Infected Lungs Upon Pulmonary Influenza Infection. Frontiers in Immunology, 2021. **Volume 12 -** 2021.

27. Huang, K., et al., BH3-only proteins target BCL-xL/MCL-1, not BAX/BAK, to initiate apoptosis. Cell Research, 2019. 29: p. 942–952.

28. Rehwinkel, J., et al., RIG-I Detects Viral Genomic RNA during Negative-Strand RNA Virus Infection. Cell, 2010. 140(3): p. 397–408.

29. Loo, Y.-M., et al., Distinct RIG-I and MDA5 Signaling by RNA Viruses in Innate Immunity. Journal of Virology, 2008. 82(1): p. 335–345.

30. Chattopadhyay, S., et al., Viral apoptosis is induced by IRF-3-mediated activation of Bax. The Embo Journal, 2010. 29(10): p. 1762–73.

31. Knowlton, J.J., T.S. Dermody, and G.H. Holm, Apoptosis induced by mammalian reovirus is beta interferon (IFN) independent and enhanced by IFN regulatory factor 3- and NF-κB-dependent expression of Noxa. Journal of Virology, 2012. 86(3): p. 1650–60.

32. Lei, Y., et al., MAVS-Mediated Apoptosis and Its Inhibition by Viral Proteins. PLOS ONE, 2009. 4(5): p. e5466.

33. Yu, C.Y., et al., The interferon stimulator mitochondrial antiviral signaling protein facilitates cell death by disrupting the mitochondrial membrane potential and by activating caspases. Journal of Virology, 2010. 84(5): p. 2421–31.

34. El Maadidi, S., et al., A Novel Mitochondrial MAVS/Caspase-8 Platform Links RNA Virus–Induced Innate Antiviral Signaling to Bax/Bak-Independent Apoptosis. The Journal of Immunology, 2014. 192(3): p. 1171–1183.

35. Zhang, Q., et al., IPS-1 plays a dual function to directly induce apoptosis in murine melanoma cells by inactivated Sendai virus. International Journal of Cancer, 2014. 134(1): p. 224–234.

36. Eitz Ferrer, P., et al., Induction of Noxa-Mediated Apoptosis by Modified Vaccinia Virus Ankara Depends on Viral Recognition by Cytosolic Helicases, Leading to IRF-3/IFN-β-Dependent Induction of Pro-Apoptotic Noxa. PLOS Pathogens, 2011. **7**(6): p. e1002083.

37. Yap, G.L.R., et al., Annexin-A1 promotes RIG-I-dependent signaling and apoptosis via regulation of the IRF3–IFNAR–STAT1–IFIT1 pathway in A549 lung epithelial cells. Cell Death & Disease, 2020. 11(6): p. 463.

38. Lallemand, C., et al., Single-stranded RNA viruses inactivate the transcriptional activity of p53 but induce NOXA-dependent apoptosis via post-translational modifications of IRF-1, IRF-3 and CREB. Oncogene, 2007. **26**(3): p. 328-338.

39. Sun, Y. and D.W. Leaman, Involvement of Noxa in Cellular Apoptotic Responses to Interferon, Double-stranded RNA, and Virus Infection Journal of Biological Chemistry, 2005. 280(16): p. 15561–15568.

