## Supplementary Figures for "RIG-I–MAVS–NOXA axis coordinates antiviral defense and apoptosis during parahenipavirus infection"

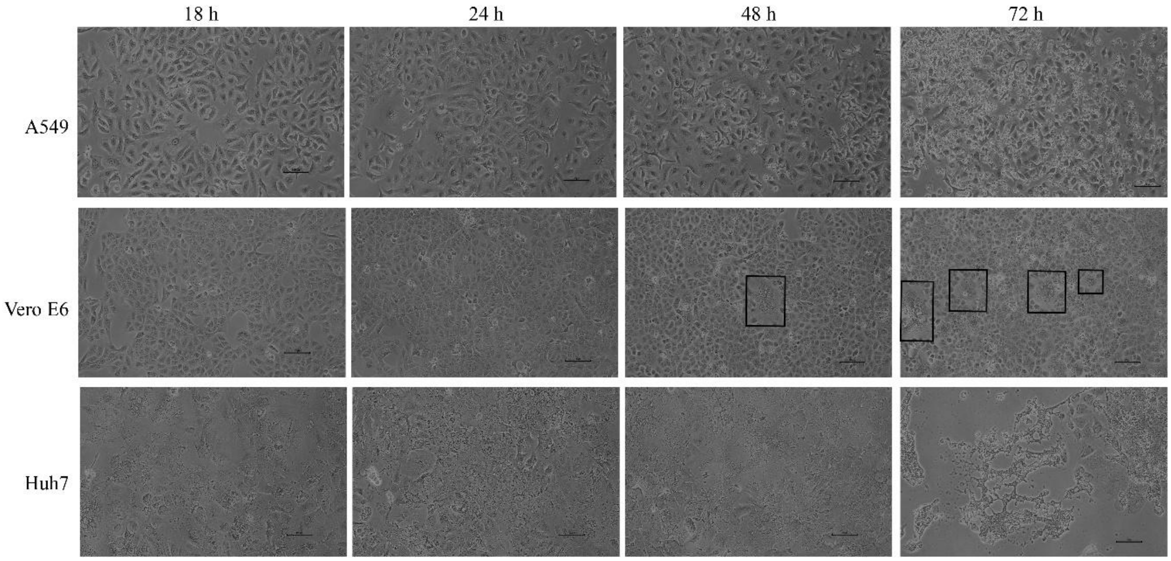


**Supplementary Figure 1. Morphological changes in mammalian cell lines following GAKV infection.** Representative brightfield images of A549, Vero E6, and Huh7 cells following GAKV infection at the indicated time points (18, 24, 48, and 72 hpi). A549 cells displayed cell rounding at 72 hpi. Vero E6 cells showed syncytia formation at both 48 h and 72 hpi (boxed region), whereas Huh7 cells exhibited marked cytopathic effects characterized by disruption of the cell monolayer and cell detachment at 72 hpi. Scale bars: 100 µm.

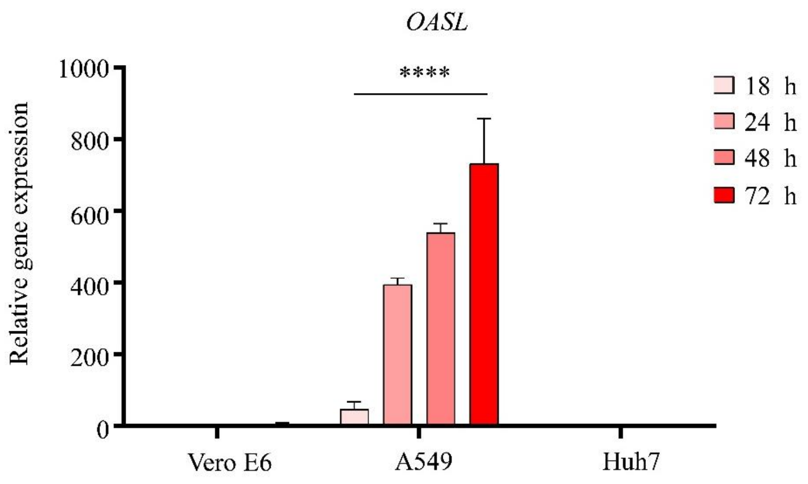


**Supplementary Figure 2. GAKV infection induces *OASL* expression in A549 cells.**

Relative *OASL* mRNA expression in A549 cells infected with GAKV (MOI = 1) at 18, 24, 48, and 72 h post-infection (hpi). Gene expression was measured by RT-qPCR and normalized to GAPDH. Data are presented as mean ± SD from three independent experiments. Statistical significance was determined using one-way ANOVA followed by Dunnett's multiple-comparison test. $P < 0.05$, $P < 0.01$, $P < 0.001$, $P < 0.0001$.

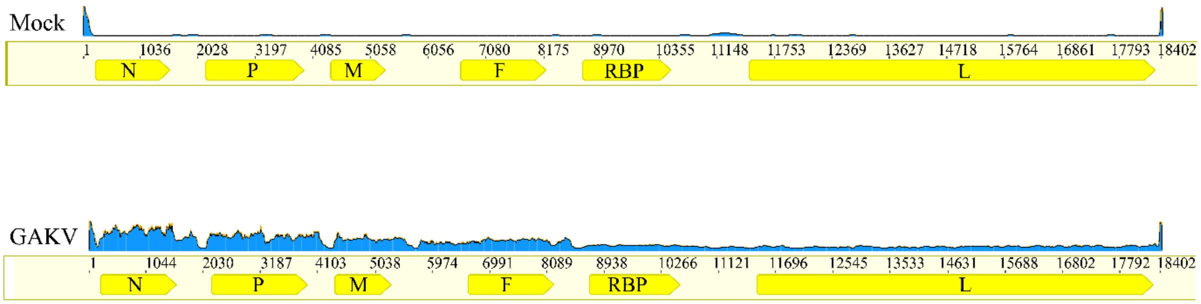


**Supplementary Figure 3. Mapping of RNA-seq reads to the GAKV genome during infection.** RNA-seq reads obtained from FASTQ files were mapped to the GAKV reference genome using Geneious Prime 2025.1.2. Coverage plots show the distribution of mapped reads across the viral genome in mock- and GAKV-infected samples at 48 hpi. Minimal background mapping was detected in mock samples, whereas infected samples showed extensive read coverage across the viral genome, including the *N*, *P*, *M*, *F*, *RBP*, and *L* genes. Increased read abundance in the GAKV-infected sample was observed toward the 3′-region of the genome, consistent with the transcriptional gradient characteristic of paramyxoviruses.

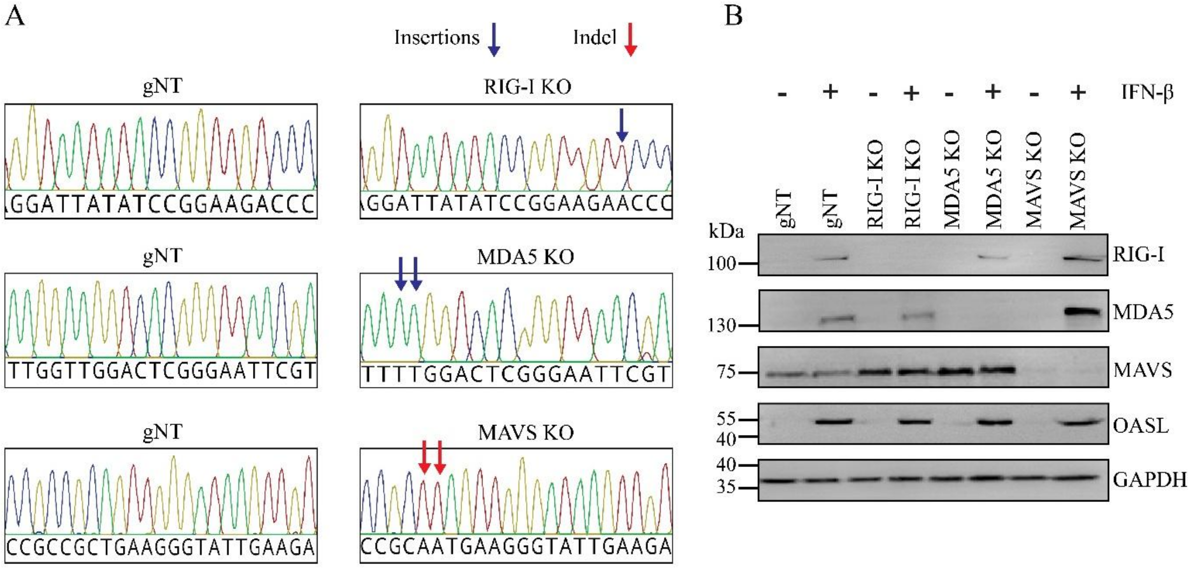


**Supplementary Figure 4. Generation and validation of RLR knockout cell lines. (A)** Sanger sequencing chromatograms showing representative sequences from control (gNT) and knockout clones for RIG-I, MDA5, and MAVS. Sequence alterations in knockout clones confirm successful genome editing at the targeted loci. **(B)** Immunoblot analysis of RLR pathway proteins in control (gNT) and knockout cell lines. Cells were treated with or without IFN-β as indicated. Protein expression of MDA5, MAVS, RIG-I, and the interferon-stimulated gene OASL was assessed. GAPDH was used as a loading control.

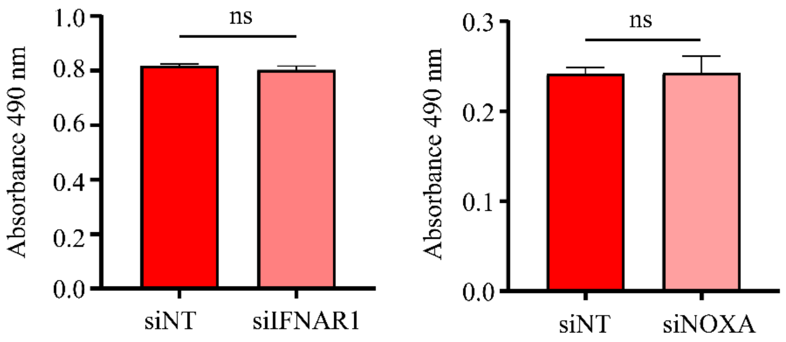


**Supplementary Figure 5. Effect of *IFNAR1* and *NOXA* knockdown on basal LDH release.** A549 cells were transfected with siNT, siIFNAR1, or siNOXA. Cell culture supernatants were collected, and LDH release was measured as an indicator of cytotoxicity. No significant differences in LDH release were observed following IFNAR1 or NOXA knockdown compared with siNT-transfected cells. Data are presented as mean ± SD from three independent experiments. Statistical analysis was performed using an unpaired Student's t-test. ns, not significant.
